# The logic of containing tumors

**DOI:** 10.1101/2020.01.22.915355

**Authors:** Yannick Viossat, Robert Noble

**Affiliations:** Ceremade, Université Paris-Dauphine, PSL, Paris, France; Department of Biosystems Science and Engineering, ETH Zurich, Basel, Switzerland; SIB Swiss Institute of Bioinformatics, Lausanne, Switzerland; Department of Evolutionary Biology and Environmental Studies, University of Zurich, Zurich, Switzerland; Department of Mathematics, City, University of London, London, UK

## Abstract

Challenging the paradigm of the maximum tolerated dose, recent studies have shown that a strategy aiming for containment, not elimination, can control tumor burden more effectively *in vitro*, in mouse models, and in the clinic. These outcomes are consistent with the hypothesis that emergence of resistance to cancer therapy may be prevented or delayed by exploiting competitive ecological interactions between drug-sensitive and resistant tumor cell subpopulations. However, although various mathematical and computational models have been proposed to explain the superiority of particular containment strategies, this evolutionary approach to cancer therapy lacks a rigorous theoretical foundation. Here we combine extensive mathematical analysis and numerical simulations to establish general conditions under which a containment strategy is expected to control tumor burden more effectively than applying the maximum tolerated dose. We show that when resistant cells are present, an idealized strategy of containing a tumor at a maximum tolerable size maximizes time to treatment failure (that is, the time at which tumor burden becomes intolerable). These results are very general and do not depend on any fitness cost of resistance. We further provide formulas for predicting the clinical benefits attributable to containment strategies in a wide range of scenarios, and we compare outcomes of theoretically optimal treatments with those of more practical protocols. Our results strengthen the rationale for clinical trials of evolutionarily-informed cancer therapy.

## Introduction

The justification for aggressive anti-cancer therapies is to maximize the probability of a cure [1,2]. This rationale disappears if a cure cannot be expected. In some if not many cases, treating aggressively can be suboptimal due to treatment toxicity and selection for resistance. A better strategy might be rather to use the *minimal* effective dose that contains the tumor subject to ensuring sufficient quality of life [3–5].

The logic of aiming for containment rather than elimination is based on evolutionary principles. At the beginning of therapy, a tumor contains cells with different sensitivities to treatment. An aggressive treatment eliminates the most sensitive cells but can enable resistant cells – freed from competing with sensitive cells for space and resources – to thrive uncontrollably. This phenomenon, called competitive release, is well understood in ecology and pest management [6–8]. By maintaining a large population of treatment-sensitive tumor cells, a containment strategy aims to exploit cell-cell competition to prevent or delay the emergence of resistance.

Various protocols in this spirit have been found to be superior to conventional therapy in experimental models [4, 9, 10], a preclinical trial [11], and a small clinical trial in metastatic castrate-resistant prostate cancer [12]. Other clinical trials are active or recruiting [13]. Yet even as empirical evidence accumulates in support of tumor containment strategies, the underlying evolutionary theory remains only imprecisely characterized in the cancer context. With the notable exception of Martin *et al.* (1992) [3,14], previous mathematical and simulation studies [4, 9, 10, 12, 15–23] have focussed on particular model formulations, specific therapeutic protocols, and typically untested assumptions about tumor growth rate, cell-cell interactions, treatment effects and resistance costs. Many previous findings are not readily generalizable because they are based on simulations, rather than mathematical analysis. Sufficient conditions for successful tumor containment have not been established. Here we address this knowledge gap by synthesizing, generalizing, and extending previous results to form a solid theoretical basis for pursuing evolutionary approaches to cancer therapy. Our work thus provides timely guidance for empirical research including the design of clinical trials.

## Model

We consider a general model with two types of tumor cells, sensitive and fully resistant, with subpopulation sizes *S*(*t*) and *R*(*t*), respectively. The total tumor population size is denoted by *N* (*t*) = *S*(*t*) + *R*(*t*), with initial value *N*_0_ = *S*_0_ + *R*_0_. Tumor dynamics are described by:

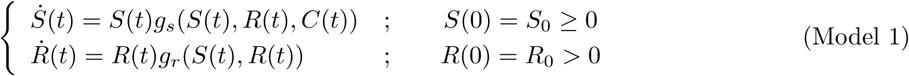

where 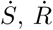 denote derivatives, and *g*_*s*_ and *g*_*r*_ are per-cell growth-rate functions; the quantity *C*(*t*) is the drug dose at time *t* (which is assumed to equate with treatment level, neglecting details of pharmacokinetics and pharmacodynamics).

For simulations, we use a Gompertzian growth model studied by Monro and Gaffney (2009) [15] (see also Martin *et al.* (1992) [3]):

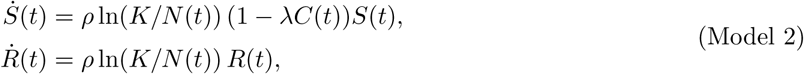

where *λ* is a sensitivity parameter, *K* is the tumor carrying capacity (the hypothetical size at which the tumor would cease to grow), and *ρ* is the baseline per-cell growth rate. We focus on this particular model in our numerical simulations to facilitate comparison with previous analysis [15], and because Gompertzian growth has been shown to describe tumor growth better than alternative models such as logistic growth [24, 25]. The parameters used for this model are those of Monro and Gaffney [15], except that we neglect mutations and backmutations after treatment initiation. They are summed up in Table 1.

**Table 1:** Parameter values. Except when otherwise specified, numerical results use the following parameter values. Model 3 is introduced later on. The initial size of the resistant subpopulation is derived through the Goldie-Coldman (1979) formula [2]: 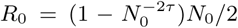, where *τ* = 10^−6^ is the mutation and backmutation rate of Monro and Gaffney [15], and *N*_0_ the initial tumor size. The value of *N*_*tol*_ is arbitrary (in log-scale, this is almost the average of *N*_0_ and *N*_*crit*_). The value of *C*_*max*_ is for consistency with Zhang *et al*.’s (2017) [12] clinical trial, where, in average, the cumulative dose given is about half the MTD (47%) Simulations were conducted in R using the deSolve package. [26].

| Parameter | Meaning | Value | Model(s) |
| --- | --- | --- | --- |
| $K$ | tumor carrying capacity | $2 \times 10^{12}$ | Model 2 |
| $K_s$ | carrying capacity of a fully susceptible tumor | $2 \times 10^{12}$ | Model 3 |
| $K_r$ | carrying capacity of a fully resistant tumor | varied | Model 3 |
| $\rho, \rho_r, \rho_s$ | baseline per-cell growth rate (per day) | 0.005928 | Models 2 and 3 |
| $\alpha$ | competition coefficient | 1 | Model 3 |
| $\beta$ | competition coefficient | varied | Model 3 |
| $\lambda$ | treatment sensitivity | 1 | Models 2 and 3 |
| $C_{max}$ | maximal instantaneous tolerated dose | 2 | Models 2 and 3 |
| $N_0$ | initial tumor size | $10^{10}$ | Models 2 and 3 |
| $R_0$ | initial resistant cell population size | $2.3 \times 10^5$ | Models 2 and 3 |
| $N_{tol}$ | tumor size corresponding to treatment failure | $7 \times 10^{10}$ | Models 2 and 3 |
| $N_{crit}$ | lethal tumor size | $5 \times 10^{11}$ | Models 2 and 3 |

*Model assumptions*. Our key assumptions are that:

- The growth rate of sensitive cells is positive in the absence of treatment, decreases as treatment dose is increased (*g*_*s*_ is strictly decreasing in *C*), and is negative at sufficiently high doses.
- Resistant cells are fully resistant (*g*_*r*_ does not depend on *C*).
- All else being equal, the larger the subpopulation of sensitive cells, the lower the growth-rate of resistant cells (*g*_*r*_ is strictly decreasing in *S*). This is a standard assumption in the adaptive therapy literature (see Supplementary material, Section 1). This might result from density-dependence (the larger the tumor, the larger its doubling time [3, 15, 27], as in the Gompertzian Model 2), frequency-dependence (the rarer resistant cells, the larger their doubling time [9,10]), a combination of those two factors [9,12,16,18,19,22], or some other form of inhibition of resistant cells by sensitive cells.
- Mutations from sensitive cells to resistants cells occurring after treatment initiation may be neglected, as well as back mutations (we checked that taking into account late random genetic mutations result in little quantitative changes in outcomes, see Supplementary Table 6).

On top of standard regularity assumptions on growth-rate functions, this is enough for our key results. Some results also require that increasing the resistant population does not increase the growth-rate of sensitive cells (*g*_*s*_ is non-increasing in *R*), excluding cooperative interactions. This ensures that the number of sensitive cells is maximized by not treating. We also use a technical assumption ensuring that the treatment level required to stabilize a tumor at a certain size increases with the frequency of resistant cells (see Supplementary material, Section 2.1). Finally, the instantaneous dose *C*(*t*) is assumed no higher than a maximal tolerated dose *C*_*max*_, but this assumption is relaxed in our idealized treatments (see below).

*Treatments*. The main treatments we consider are the following:

- *Maximal Tolerated Dose (MTD)*: *C*(*t*) = *C*_*max*_ throughout.
- *Containment at the initial tumor size N*_0_: this treatment continuously adjusts the dose to maintain total tumor size at *N* (*t*) = *N*_0_ as long as possible with a dose *C*(*t*) *≤ C*_*max*_, then treats at *C*_*max*_. Mathematically, the stabilizing dose is found by solving the equation 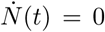. In the Gompertzian Model 2, this leads to *C*(*t*) = *N* (*t*)*/λS*(*t*). The dose administered is the minimum of this stabilizing dose and of *C*_*max*_. In practice, containment would only be approximative, and the appropriate dose would be found by regular monitoring of the patient and dose adjustments. This would not require to differentiate between sensitive and resistant cells. Possible protocols are discussed on page 12 and in Supplementary material, Section 6.
- *Containment at some other threshold size N* ^*∗*^: this treatment does not treat until tumor size reaches *N* ^*∗*^ (if *N* ^*∗*^ *≥ N*_0_), or treats at the maximal tolerated dose until tumor size is reduced to *N* ^*∗*^ (if *N* ^*∗*^ *< N*_0_), and then contains the tumor at this threshold as above.

To reveal the logic of containment as clearly as possible, we also consider idealized versions of these treatments, with no constraint on the maximum instantaneous dose (so that the sensitive population can be reduced instantly to any desired size). These idealized treatments, though biologically unrealistic, help reveal the basic logic of containment and provide reference points largely independent of model details. In the idealized form of maximum tolerated dose treatment (*ideal MTD*), the sensitive population is instantly eliminated (so that *S*(*t*) = 0 for all *t >* 0). This is called “aggressive treatment” or “elimination of sensitive cells” by Hansen *et al.* (2017, 2019) [18] [28] and Hansen and Read (2020) [27]. We may think of this as a treatment inducing an infinite kill-rate. *Ideal containment at the initial tumor size* maintains the tumor at its initial size as long as some sensitive cells remain. The tumor is then fully resistant, hence its later growth independent of the treatment. *Ideal containment at some other threshold N* ^*∗*^ lets the tumor grow to *N* ^*∗*^ (or instantly reduces tumor size to *N* ^*∗*^, if *N* ^*∗*^ *< N*_0_), then stabilizes tumor size at this threshold as long as some sensitive cells remain. Containment in the sense of Hansen *et al.* (2017) [18], from which we borrow this vocabulary, corresponds to our ideal containment treatment, except that we do not allow for an instantaneous increase in tumor size.

Containment and MTD treatments are illustrated in Figure 1. We also consider other possibilities such as *constant dose or delayed constant dose treatments*, studied by Monro and Gaffney (2009) [15]; *intermittent containment* (Fig. 1g), where tumor size is maintained between a high and a low threshold, as in Zhang *et al* (2017) [12]; and forms of *metronomic therapy*, where treatment is turned on and off at predefined times.

**Figure 1:**
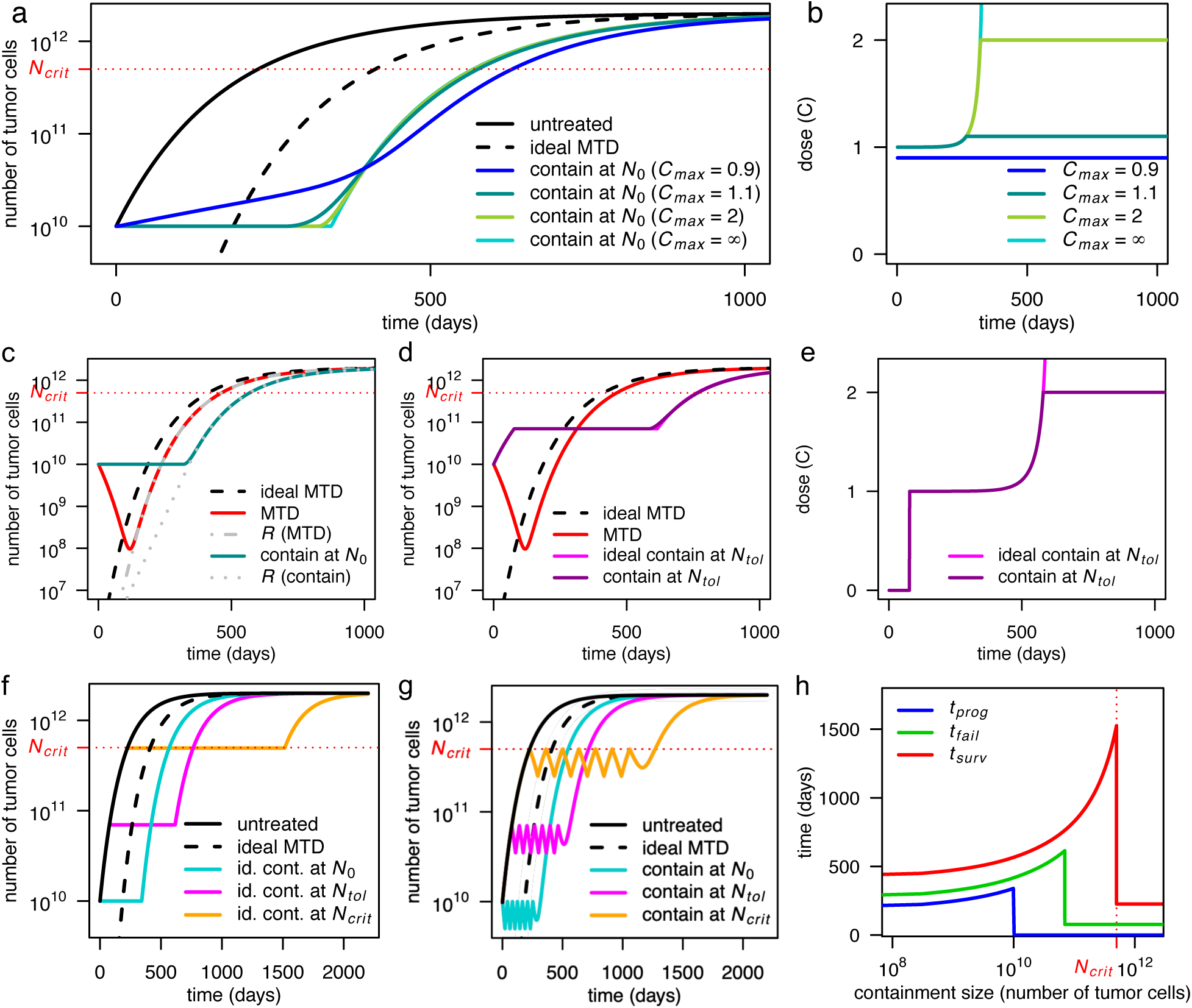
Illustration of containment and MTD treatments in Model 2. **a**, Tumor size under no treatment (black), ideal MTD (dashed), and containment at the initial size for various values of the maximum tolerated dose *C*_*max*_. The case *C*_*max*_ = *∞* (light blue) corresponds to ideal containment. The patient is assumed to die shortly after tumor size becomes greater than *N*_*crit*_. **b**, Drug dose under the containment treatments of panel a. If *C*_*max*_ *<* 1, the tumor cannot be stabilized and containment boils down to MTD. **c**, Tumor size under MTD, ideal MTD, and containment at the initial size, and resistant population size under MTD and containment. **d**, Tumor size under MTD, containment at the maximum tolerable size, and their idealized counterparts. **e**, Drug dose under containment and ideal containment at the maximum tolerable size, as represented in panel d. **f**, Tumor size under no treatment, ideal MTD, and ideal containment at three different tumor sizes. **g**, Tumor size under no treatment, ideal MTD, and intermittent containment between *N*_*max*_ and *N*_*min*_ = *N*_*max*_*/*2 for three different values of *N*_*max*_. **h**, Times to progression (blue), treatment failure (green), and survival time (red) under ideal containment at a threshold size varied from *R*_0_ to *N*_*crit*_ (ideal containment at *R*_0_ is equivalent to ideal MTD). The time until the tumor exceeds a certain size is maximized by ideal containment at that size. Exact formulas for idealized treatments are in Supplementary material, Section 3.1.

*Outcomes.* Our three main outcomes are :

- *Time to progression:* defined here as the time until the tumor exceeds its initial size, *N*_0_. The RECIST criterion is that progression occurs when tumor size is 20% larger than at treatment initiation. This 20% buffer makes sense in medical practice, due to imperfect monitoring of the tumor and imperfect forecast of treatment’s effect. In our mathematical models however, this buffer is not needed, and would only obscure the analysis, so we use a more basic definition.
- *Time to treatment failure:* until the tumor exceeds a threshold size determined by the physician and patient, *N*_*tol*_, which we call the maximal tolerable size. This may be thought of as the maximal tumor size at which the tumor is not quickly life threatening, based on physician’s expertise, and does not result in too severe side effects for the patient. Due to this second requirement, the maximal tolerable size would only be revealed during treatment. To fix ideas, we assume that it is higher than the initial tumor burden, *N*_0_. The case where it is lower is studied in Supplementary Material.
- *Survival time:* until the tumor reaches an hypothetical lethal size, *N*_*crit*_, after which the patients is assumed to die quickly. This lethal tumor burden is also patient specific.

## Results

### When is containment optimal?

The optimal treatment strategy depends on the clinical objective. If the emphasis is on rapidly reducing tumor burden then the maximum tolerated dose (MTD) is clearly superior to containment. However, if the aim is to maximize time to progression, then our formal mathematical analysis proves that containment is likely to be optimal, or at least close to optimal, in a broad range of cases.

To see why, consider a tumor containing sensitive and fully resistant cells. The growth rates of these two subpopulations are expected to depend on the subpopulation sizes, and the growth rate of sensitive cells will also vary with the treatment dose. Furthermore, if resource competition is the dominant ecological interaction between subpopulations then it is reasonable to assume that, all else being equal, the larger the sensitive population, the lower the growth rate of the resistant population. To the best of our knowledge, this latter assumption holds for all proposed mathematical models with two cell types in which the impact of mutations after treatment initiation can be neglected (see Section 1 of Supplementary material for a review of previous studies).

If the objective is to maximize time to progression then, under the above general assumptions, we find that the best possible treatment is the containment strategy that precisely maintains the original tumor burden for as long as some sensitive cells remain, which we called ideal containment. Moreover, among treatment strategies that eventually eliminate the sensitive population, we find that the worst option is to maximise the cell kill rate, that is, the ideal MTD treatment. Instead of maximizing time to progression, an alternative objective is to maximize the time until tumor burden exceeds a certain threshold. In this case, the optimal treatment maintains the tumor at precisely this threshold size.

The intuitive explanation is that, whereas we can always reduce the sensitive population by using a sufficiently aggressive treatment, the only way to impair the growth of resistant cells is to exploit competition with sensitive cells. By assumption, this ecological form of control is most effective when the sensitive population is as high as can be permitted. Conversely, competition is least effective when the sensitive population is smallest; that is, under MTD.

Which containment strategy works best depends on the objective (Fig. 1h). Time to progression is maximized by ideal containment at the initial size; time to treatment failure, by ideal containment at the maximal tolerable size. In theory, survival time would be maximized by ideal containment just below the lethal tumor size. Attempting this would however be extremely dangerous, both due to adverse effect on patient’s quality of life and because too optimistic a guess of the lethal burden would lead to quick patient’s death.

### Mathematical intuition

Formal mathematical proofs of all these results in Model 1 can be found in Supplementary material, Section 2. They are based on a differential equation tool called the comparison principle (a variant of Gronwall’s lemma), but the basic argument is simple (see also [18]): between time *t* and *t* + *dt* (where *dt* is a small time increment), the resistant population increases from *R*(*t*) to *R*(*t* + *dt*) ≃ *R*(*t*) + *R*^*t*^(*t*)*dt*, hence by a quantity

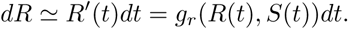

So, if we fix a resistant population size *R*_1_ and a small size increment *dR*, the time it takes for the resistant population size to grow from *R*_1_ to *R*_1_ + *dR* is roughly:

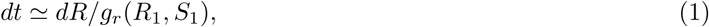

where *S*_1_ is the sensitive population size when *R* = *R*_1_. Assuming *R*_0_ *≤ R*_1_ *≤ N*_0_, under ideal containment at the initial size, *S*_1_ +*R*_1_ = *N*_0_, so *S*_1_ = *N*_0_ −*R*_1_. Before progression, under any other treatment, *S*_1_ *≤ N*_0_ −*R*_1_. By assumption, the larger the sensitive population, the lower the resistant population growth rate, hence the higher the duration *dt* in (1); it follows that the time it takes for the resistant population to grow from *R*_1_ to *R*_1_ + *dR* is maximized by ideal containment (and minimized by ideal MTD, since then *S*_1_ = 0). Iterating this argument shows that the resistant population *R*_*idcont*_(*t*) under ideal containment at the initial size will be smaller than the resistant population *R*(*t*) under any alternative treatment, at least as long as none of these treatments led to progression. Since under ideal containment at the initial size, progression occurs when *R*_*idcont*_(*t*) = *N*_0_, this implies that progression occurs later than under any other treatment. Other results require more sophisticated arguments, but the intuition is similar.

### Characterizing the intensity of competition between tumor cells

Although we find that the superiority of ideal containment is qualitatively very robust, the predicted magnitude of clinical benefits depends on biological assumptions and parameter values. A critical factor is the strength of competition between treatment-sensitive and resistant cells. One biologically plausible hypothesis is that the growth rate of resistant cells primarily depends on their abundance relative to sensitive cells [9,10]. In ecological parlance, this means that the fitness of resistant cells is *frequency-dependent*. If this assumption holds then the most important parameter in determining outcomes is the relative fitness of resistant cells when rare [9]. In other words, what matters is how rapidly resistant cells proliferate, relative to sensitive cells, while resistant cells make up only a tiny fraction of the tumor. The smaller this parameter value, the greater the predicted clinical gains from containment, relative to MTD.

An alternative, equally plausible hypothesis is that resistant cells fitness is primarily *density-dependent*, such that the per-cell growth rate decreases as the total tumor burden increases [3, 12, 14, 15, 18]. What matters most in this case is the strength of density dependence. In the widely-used Gompertzian model of tumor growth, the per-cell growth rate decreases relatively rapidly with increasing tumor size, leading to a strong competition effect and substantial clinical gains for containment versus aggressive treatment. Mathematical models that describe weaker competition, such as the logistic growth model, predict smaller clinical gains [3], whereas those that describe stronger competition, such as the von Bertalanffy growth model, predict larger gains (Fig. 2c, Supplementary Fiig. 1; Supplementary material, Section 4.1.1).

**Figure 2:**
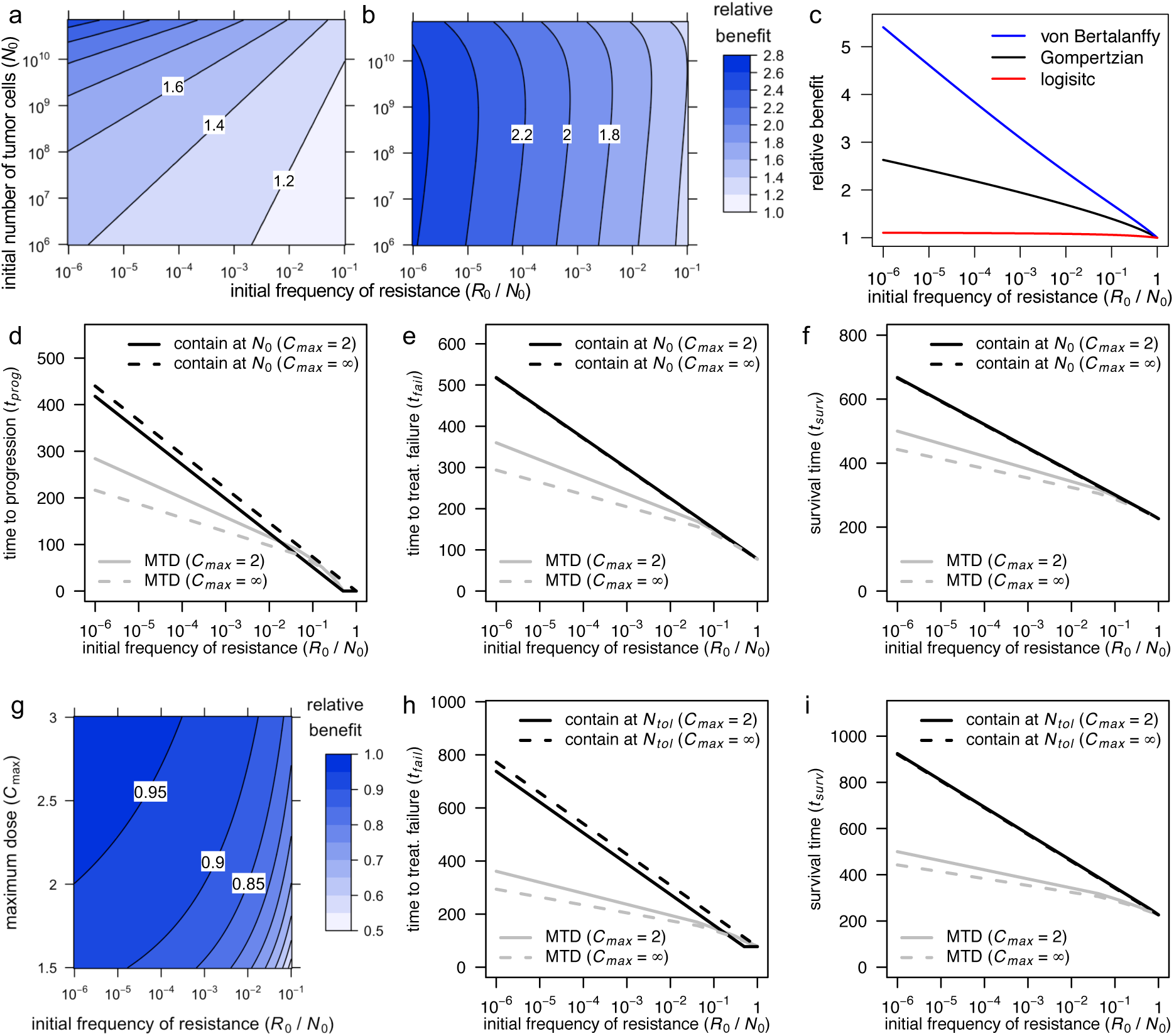
Comparison of clinical benefits of containment and MTD treatments in Model 2. **a**, Relative benefit, in terms of time to progression, for ideal containment at size *N*_0_ versus ideal MTD (that is, ratio *t*_*prog*_ (*idContN*_0_)*/t*_*prog*_ (*idMT D*)), as a function of initial tumor size and frequency of resistant cells. **b**, Relative benefit, in terms of time to treatment failure, for ideal containment at size *N*_*tol*_ versus ideal MTD (that is, ratio *t*_*fail*_(*idContN*_*tol*_)*/t*_*fail*_(*idMT D*)), as a function of initial tumor size and frequency of resistant cells. **c**, Relative benefit, in terms of time to treatment failure, for ideal containment at size *N*_*tol*_ versus ideal MTD, for a Gompertzian growth model (black curve; Model 2), a logistic growth model (red) and a von Bertalanffy growth model (blue). Parameter values for the Gompertzian growth model are as in Table 1. Parameter values of the other two models are chosen so that untreated tumor growth curves are similar for tumor sizes between *N*_0_ and *N*_*crit*_ (the lethal size). See Supplementary Fiigure 1 for details. **d, e, f**, Time to progression (panel d), to treatment failure (panel e), and survival time (panel f) versus initial frequency of resistance. Outcomes are shown for MTD treatment and containment at *N*_0_, both in the ideal case (*C*_*max*_ = *∞*) and subject to *C*_*max*_ = 2. **g**, Relative benefit, in terms of time to treatment failure, for containment versus ideal containment (at size *N*_*tol*_), as a function of maximum dose threshold (*C*_*max*_) and initial frequency of resistant cells (formulas are in Supplementary material, Section 3.3). Contour lines are at intervals of 0.05. **h, i**, Time to treatment failure (panel h), and survival time (panel i) versus initial frequency of resistance. Outcomes are shown for MTD treatment and containment at *N*_*tol*_, both in the ideal case (*C*_*max*_ = *∞*) and subject to *C*_*max*_ = 2.

Important differences between model predictions underscore the need to advance understanding of the ecological interactions that govern intra-tumor dynamics [29], which remain only poorly characterized. Nevertheless, we find that what matters is not so much whether a mathematical model assumes frequency- or density-dependent fitness, but rather whether the model accurately describes the strength of this dependence.

### Other important biological parameters

To illustrate how biological parameter values are predicted to influence clinical outcomes, consider Model 2. Recall that the ideal MTD treatment depicted in Fig. 1a (dashed line) is an idealized version of MTD which instantly eliminates sensitive cells [18] [28]. The clinical gain from this regimen, compared to no treatment, is the time it takes for the tumor to grow from *R*_0_ (the initial resistant population) back to its initial size *N*_0_; that is, the time taken for the resistant population to increase by a factor of *N*_0_*/R*_0_ after being freed from competition with sensitive cells.

In the case of ideal containment, the clinical gain is instead the duration of the stabilization phase. This is equal to the time taken for the resistant population to increase by a factor of *N*_0_*/R*_0_ while the tumor is maintained at the stabilization size. Because competition with sensitive cells impedes the growth of resistant cells, the gain from ideal containment is invariably greater than the gain from ideal MTD. Moreover, a larger stabilization size results in a longer stabilization phase and hence improved survival (Fig. 1f).

Simple mathematical expressions may be derived to quantify the effects of containment and MTD strategies in various density-dependent scenarios, and in some frequency-dependent ones (see Supplementary material, Section 3). This enables us to examine the impact of varying any parameter on time to progression, time to treatment failure, and survival time. Recall that these three outcomes are defined, respectively, as the times until the tumor becomes larger than *N*_0_ (the size at treatment initiation), *N*_*tol*_ (a hypothetical maximum tolerable size) and *N*_*crit*_ (the hypothetical lethal tumor size). For idealized treatments, these outcomes are independent of the treatment’s mode of action (for example, whether it results in a log kill rate, a Norton-Simon kill rate proportional to the net growth rate of an untreated tumor [1], or some other effect).

For Model 2, the times to progression under ideal containment at the initial size and ideal MTD are

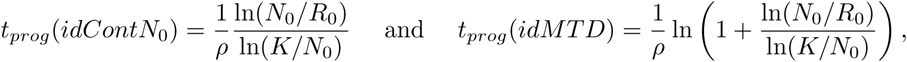

respectively, where ln is the natural logarithm. In terms of time to progression, the *absolute* clinical benefit of ideal containment over ideal MTD is the difference between these numbers; the *relative* benefit (or *fold change in progression time* in Hansen and Read [27]) is the ratio

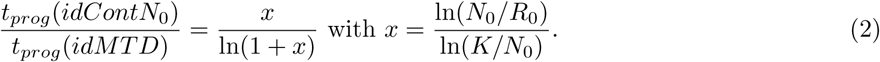

These formulas reveal the importance of three patient-specific factors: the baseline growth-rate, *ρ*; the initial frequency of resistant cells, *R*_0_*/N*_0_; and the initial tumor size compared to the carrying capacity, *N*_0_*/K*.

For idealized treatments, decreasing the growth rate parameter (*ρ*) has no effect on the *relative* clinical benefits of containment, but, by slowing the dynamics, leads to higher *absolute* benefits. Instead decreasing the initial frequency of resistant cells (*R*_0_*/N*_0_) increases both absolute and relative clinical gains of containment versus MTD. This is in part because aggressive treatments are especially suboptimal when resistance is very rare, as they then cause a drastic reduction in tumor size, which permits rapid expansion of the resistant population. Lastly, a higher value of ratio *N*_0_*/K* implies more intense competition at the initial tumor size. This increases both absolute and relative benefits of containment at the initial size. Fig. 2 illustrate some of these effects for Model 2. The impact of a large initial tumor size on relative benefits of containment at the maximal tolerable size is more complex (Fig. 2b; Supplementary material, Section 4.1.2).

### Practical treatment strategies can be close to optimal

For simplicity in the above mathematical analysis, we assumed no restriction on maximum dose, which permits the ideal containment strategy of maintaining the tumor precisely at a target size until it becomes fully resistant. In reality, toxicity constraints typically impose a maximum instantaneous dose *C*_*max*_. Figs. 1a, 1b, 1d and 1e compare tumor dynamics and doses under ideal containment and under containment strategies. In the latter case, the stabilization phase is shorter because it finishes before all sensitive cells have been removed, which results in shorter times to progression or treatment failure (Figs. 2d, 2g, 2h). However, as long as *C*_*max*_ is substantially higher than the dose allowing to stabilize a fully sensitive tumor, the two treatments are quite close and result in similar survival times (Figs. 1a, 1d, Figs. 2f, 2h). Differences between ideal and non-ideal containment outcomes are discussed in greater detail in Supplementary material, Section 4.2.

An additional consideration is that a continuous containment strategy requires continuous monitoring of tumor size, which is typically infeasible. More practical protocols include intermittent containment, constant dose therapy and metronomic therapy.

#### Intermittent containment

The question of whether it is better to implement containment via a continuous low dose or an intermittent high dose treatment has yet to be settled. Both strategies worked well in mice [11]. Although Zhang *et al.* (2017) [12] obtained highly promising clinical results from intermittent high dose treatment, it is plausible that a continuous low dose treatment would have performed even better (as, if anything, seems to be the case in mice [11], although the evidence is too scarce to be conclusive). Mathematical models that account for cell-cycle dynamics, pharmacodynamics, and drug-induced resistance may be able to predict the optimality of a specific intermittent treatment, provided they can be precisely parameterized. In our simple setting, however, higher tumor burden implies slower growth of resistance, and hence exact containment at an upper threshold *N*_*max*_ is better than containment between upper and lower bounds *N*_*max*_ and *N*_*min*_. Nevertheless, as is apparent from comparing Figs. 1g and 1h, and from the theoretical analysis in the Supplementary material (Section 2.3, Propositions 5 and 6; Section 3.1.5; Section 3.3, Supplementary Table 4), the difference between the two types of protocol is small provided that *N*_*min*_ is a large fraction of *N*_*max*_. Indeed, decreasing tumor burden from *N*_*max*_ to *N*_*min*_ then only slightly increases the growth rate of resistant cells. Intermittent containment as in the clinical trial of Zhang *et al.* (2017) [12], which may be the only practical possibility, therefore appears to be a sound implementation of containment, as long as the lower threshold is not too low.

#### Constant dose

To maximize time to progression in Model 2, the optimal constant dose is slightly higher than *C* = 1*/λ* (which corresponds to *C* = 1 in Fig. 3a). The constant dose *C* = 1*/λ* stabilizes the sensitive population size, whereas containment uses the evolving dose *C* = *N/λS* = 1*/λ* + *R/λS* to stabilize tumor size. According to our definition, the former approach leads to immediate progression because it allows the overall tumor size to increase from the start of treatment. However, provided that resistant cells are initially rare, the dose *C* = 1*/λ* maintains tumor size close to the initial size for nearly as long as under containment (Figs. 3a, 3c). Differences that emerge after resistant cells become abundant are relatively unimportant. Thus, for a given patient, the dose *C* = 1*/λ* is expected to lead to similar outcomes as containment at the initial size. Similarly, delaying treatment until the tumor size reaches *N*_*tol*_ and then applying dose *C* = 1*/λ* has similar outcomes as containment at the maximum tolerable size (Figs. 3b, 3e). Table 2 gives examples of times to progression, times to treatment failure, and survival times for various constant doses and other treatments. Note that the constant dose that maximizes time to progression is slightly higher than 1*/λ*, whereas the non-delayed constant dose that maximizes survival time is lower than 1*/λ* (Fig. 3c, Supplementary Table 6, Supplementary Fiig. 4). Constant dose treatments may lead to higher survival time than containment at the initial size (Fig. 3a, Supplementary Fiig. 4) but to the cost of quicker progression, and they always lead to lower survival time than containment at sufficiently higher sizes.

**Table 2:** Time to progression, time to treatment failure, and survival time for Model 2. The constant dose or delayed constant doses *C* = 1.09 and *C* = 1.07 maximize *t*_*prog*_ and *t*_*fail*_, respectively, among all constant dose or delayed constant dose treatments. Times are measured in days.

| Treatment | $t_{prog}$ | $t_{fail}$ | $t_{surv}$ |
| --- | --- | --- | --- |
| No treatment | 0 | 77 | 226 |
| Ideal MTD | 186 | 263 | 412 |
| MTD ($C_{max} = 2$) | 236 | 314 | 463 |
| $C = 1.09$ | 303 | 397 | 549 |
| Containment at $N_0$ ($C_{max} = 2$) | 318 | 418 | 568 |
| Ideal containment at $N_0$ | 340 | 417 | 566 |
| $C = 1.07$ from $N = N_{tol}$ | 0 | 543 | 731 |
| Containment at $N_{tol}$ ($C_{max} = 2$) | 0 | 580 | 767 |
| Ideal containment at $N_{tol}$ | 0 | 615 | 764 |

**Figure 3:**
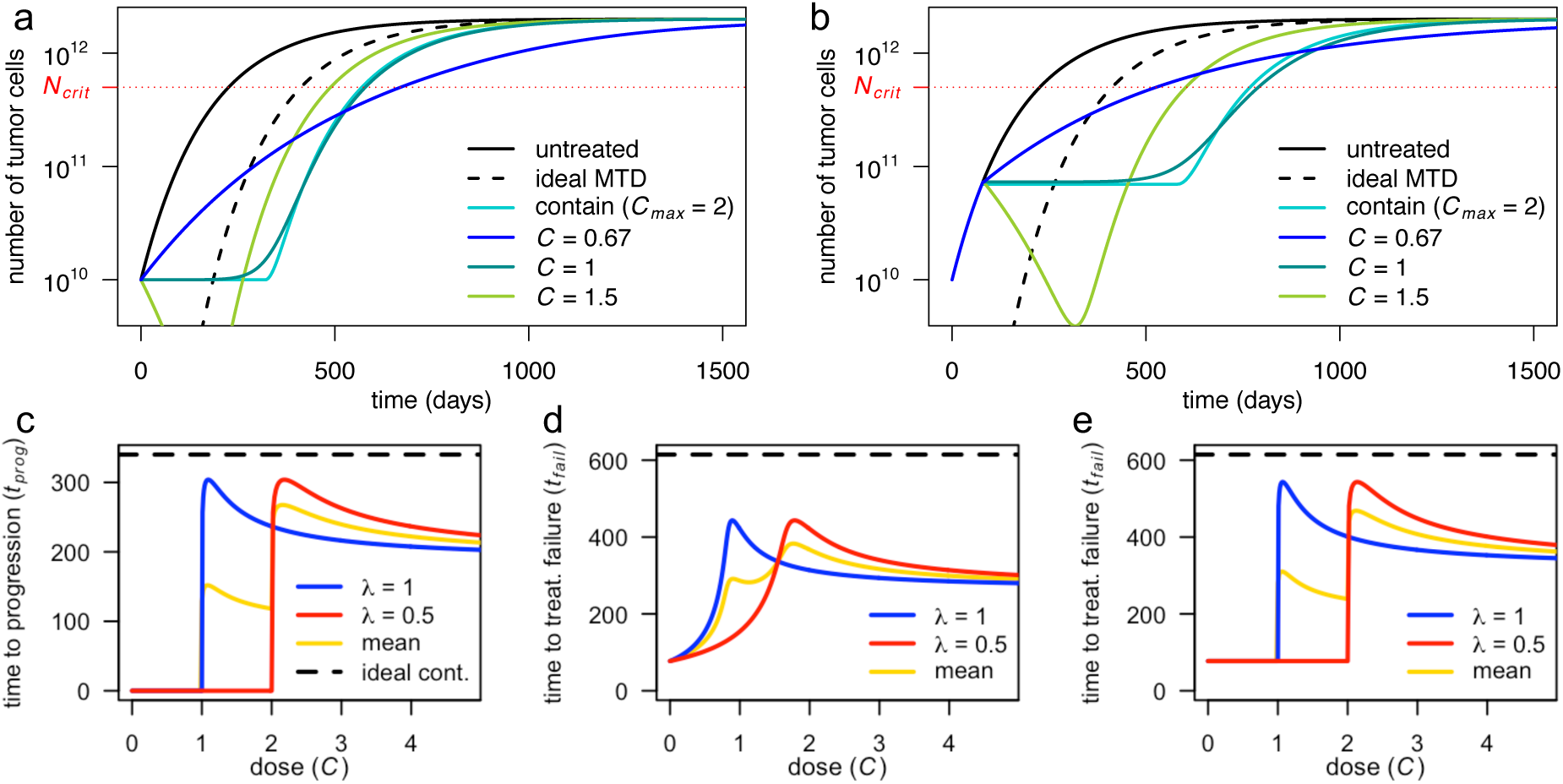
Constant dose and delayed constant dose treatments in Model 2. **a**, Tumor size for various constant dose treatments compared to containment at the initial size (subject to *C*_*max*_ = 2) and ideal MTD. **b**, Tumor size for various delayed constant dose treatments (the dose is applied continuously from the first time when *N* = *N*_*tol*_) compared to containment at *N*_*tol*_ (subject to *C*_*max*_ = 2) and ideal MTD. Until *N* = *N*_*tol*_, all curves are the same, except ideal MTD. **c**, Times to progression for two patients whose tumors differ in treatment sensitivity (parameter *λ*) under constant dose treatments, as a function of the dose. The yellow line is the mean of the two patient outcomes and the dashed line is the time to treatment failure under ideal containment at *N*_0_ (which is the same for both patients, and the maximal time to progression). **d**, Times to treatment failure for two patients whose tumors differ in treatment sensitivity under constant dose treatments, as a function of the dose. The yellow line is the mean of the two patient outcomes and the dashed line is the time to treatment failure under ideal containment at *N*_*tol*_ (which is the same for both patients, and the maximal time to treatment failure). **e**, Times to treatment failure for two patients whose tumors differ in treatment sensitivity under delayed constant dose treatment (the dose starts to be applied when *N* = *N*_*tol*_ for the first time). The yellow line is the mean of the two patient outcomes and the dashed line is the time to treatment failure under ideal containment at *N*_*tol*_ (which is the same for both patients, and the maximal time to treatment failure).

#### Adaptive treatments may be close to optimal for all patients

A problem with constant dose therapy is that the parameters that determine the best dose for a particular patient are typically unknown. Giving slightly too little or too much treatment can be far from optimal (blue and red curves in Figs. 3c, 3d, 3e). Any constant dose that works relatively well for some patients will inevitably be suboptimal for others, and the constant dose that gives the best average result for a cohort of patients will typically be further from containment than the best constant dose for a single patient (Figs. 3c, 3d, 3e; Supplementary material, Section 4.6). By contrast, a containment strategy will be close to optimal for every patient because it entails continuously adjusting the dose as a function of patient response, without requiring any parameter to be known in advance (except that the tolerable tumor burden *N*_*tol*_ must be chosen by the physician or revealed during treatment). Similarly (in the absence of an initial induction phase where treatment is given at MTD, which would trigger competitive release), conventional metronomic therapy – in which low doses are given at regular, predefined intervals – may look similar to intermittent containment. However, intermittent containment (a particular form of adaptive therapy [12]) has the important additional benefit of adapting doses to the evolution of the tumor and to patient-specific parameters, without knowing these parameters in advance [30].

### Fitness costs of resistance are helpful but not essential

Previous studies have asserted or assumed that a necessary condition for effective tumor containment is that treatment resistance incurs a cellular fitness penalty. This is especially the case in the literature on adaptive therapy (for example, [4, 5, 10, 11, 30–33]). As noted in a recent review article [5], “the theory behind adaptive therapy focuses on the phenotypic costs of the molecular mechanism(s) of resistance.” Importantly, none of our qualitative results depends on a fitness cost of resistance. Indeed, we find that containment can be an optimal and highly effective strategy even if resistant cells are fitter than treatment-sensitive cells in the absence of therapy.

Yet, although not required for containment to improve on aggressive treatment, fitness costs of resistance may increase clinical gains. The precise effect depends on whether the cost of resistance is constant or is higher in the presence of sensitive cells [18]. To examine this issue, let us consider the following model:

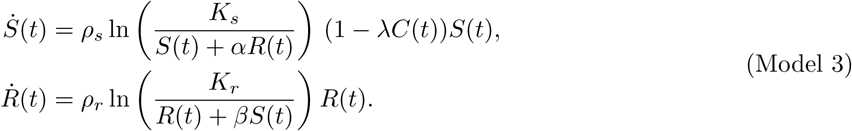

Here, the baseline growth rates *ρ*_*s*_, *ρ*_*r*_ and the carrying capacities *K*_*s*_, *K*_*r*_ are specific to sensitive and resistant cells, respectively. In the denominators, total tumor size has been replaced by a weighted sum of the resistant and sensitive population sizes, as is commonly assumed in ecological models. The higher the competition coefficient *β*, the greater the impact of sensitive cells on resistant cells. If *β* = 1, then resistant cells are affected equally by all cells and *R* + *βS* = *N*, as in Model 2.

In Model 3, a resistance cost may correspond to:

- a reduction in growth rate, independent of competition intensity (low *ρ*_*r*_);
- a general inability to compete with other cells (low *K*_*r*_);
- a specific inability to compete with sensitive cells (high *β*).

All such costs increase clinical gains of any treatment by slowing the growth of resistant cells. Moreover, resistance costs reduce the expected initial fraction of resistant cells, which is good for all treatments, but especially good for containment.

Nevertheless, for a given initial fraction of resistant cells, only some types of resistance cost increase relative clinical gains of ideal containment over ideal MTD. For instance, halving *ρ*_*r*_ doubles times to progression under both treatments, so the absolute benefit of containment doubles, but the relative benefit is unchanged. In a model that accounts for mutation from sensitive to resistant, lowering *ρ*_*r*_ may even decrease the relative benefit of containment [18]. In contrast, lowering *K*_*r*_ or increasing *β* increases both absolute and relative clinical gains, because it harms resistant cells proportionally more in the presence of sensitive cells. Some of these effects are illustrated in Figure 4 (see also complementary analyses in [18, 34]).

**Figure 4:**
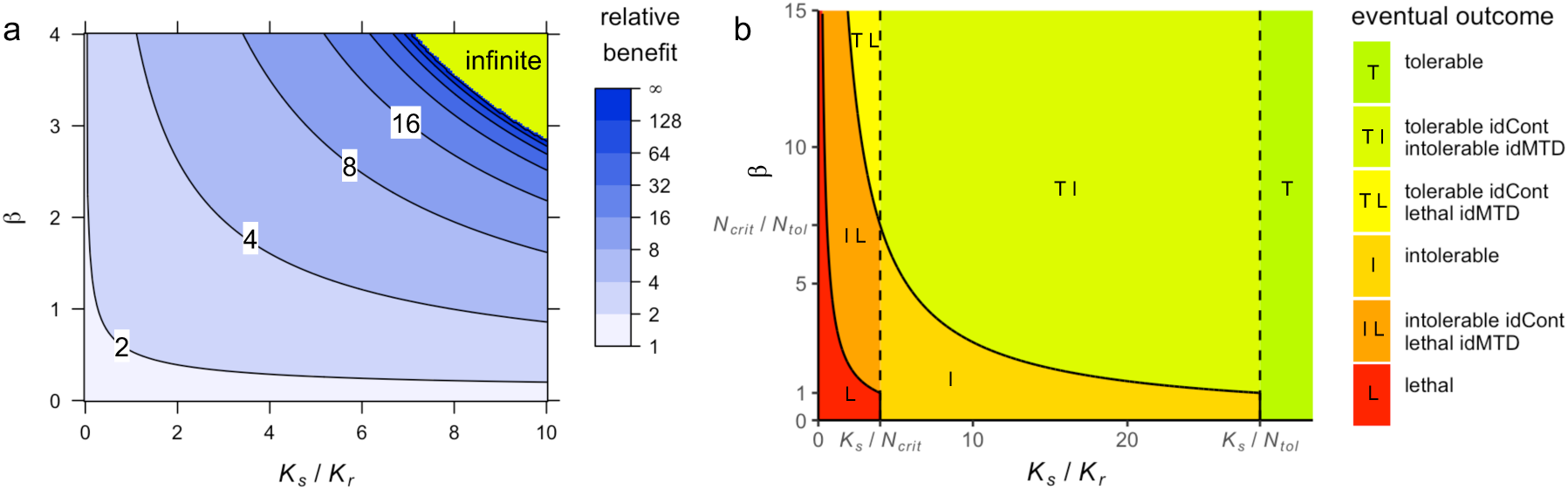
Consequences of costs of resistance in Model 3. **a**, Relative benefit, in terms of time to treatment failure, for ideal containment (at size *N*_*tol*_) versus ideal MTD, for varied values of *K*_*r*_ and *β*. This figure is based on approximate formulas that are highly accurate for the selected parameter values (see Supplementary material, Section 5.2). Supplementary Fiig. 5 shows an alternative version of this plot based on simulations. Contour lines are at powers of 2. **b**, Eventual outcomes of ideal containment (idCont) and ideal MTD (idMTD) treatment strategies, based on exact formulas (see Supplementary material, Section 5.1). The “infinite” region in panel a corresponds to the “TI” region in panel b. Fixed parameter values are as in Table 1.

### What if all tumor cells are partially sensitive to treatment?

If resistant cells retain some sensitivity to treatment then the basic logic changes in two ways. First, if resistant cells are sufficiently sensitive, then MTD can cure the tumor. This is not a case that concerns us, since our goal is to find alternative treatments when MTD is expected to fail. Second, even if a cure is impossible, there are now two ways to fight resistant cells: treating at low dose (to maintain competition with sensitive cells) or aggressively (to exploit partial sensitivity). Since competition with sensitive cells weakens as the sensitive population is depleted, treatment failure can be delayed by switching from a containment strategy to MTD at an appropriate time before treatment failure, but at the cost of increased toxicity. Whether the gain from switching to MTD is typically small or substantial remains to be investigated but, in general, the difference in outcomes for containment versus MTD is smaller when all cells are partially sensitive to treatment. If resistant cells are sufficiently sensitive then MTD may even be superior to pure containment.

If resistant cell frequency and sensitivity are unknown then we face a conundrum. Should we treat at high dose after low dose treatment failure? If the tumor is already fully resistant then any further treatment will incur needless toxicity. If resistant cells are fully resistant but some sensitive cells remain then it might be better to maintain a low dose. But if resistant cells retain some sensitivity then treating at high dose after initial treatment failure may be the best option, subject to treatment toxicity. To make the best choice, clinicians will require new methods for assessing tumor composition and sensitivity during therapy. Determining optimal strategies in the case of partial or unknown treatment sensitivity is an important topic for future theoretical research. In particular, when it may be proved that it is optimal to first contain the tumor and then switch to MTD, clinically implementable methods to determine a close to optimal switching time should be developed.

### When can the tumor be contained forever?

In Model 3, unless a fully sensitive or fully resistant tumor is intrinsically benign (*K*_*s*_ *< N*_*tol*_ or *K*_*r*_ *< N*_*tol*_, respectively), indefinite containment under the maximum tolerable size requires two conditions: first, resistant cells are harmed more from competition with sensitive cells than from competition with other resistant cells (*β >* 1); second, the resistant population would decline in an almost fully sensitive tumor of threshold size *N*_*tol*_.

The latter condition is equivalent to *K*_*r*_ *< βN*_*tol*_. Since the resistant population’s carrying capacity is likely to be significantly larger than the threshold tumor size, this condition typically requires a large competition coefficient *β*. Therefore, at least in this model, indefinite containment is possible only if sensitive cells greatly impair the fitness of resistant cells (green region of Figure 4a; green and yellow regions of Figure 4b). These results are derived in Supplementary material, Section 5.1.

### Protocols for containment at the initial size

Previous studies suggested variants of downward or upward titration strategies, where the dose is gradually decreased or increased until an appropriate stabilizing dose is found [4, 11, 17, 35]. This may be combined with treatment vacations. Although titration methods have the advantage of being conceptually simple, they could be slow in determining an approximately stabilizing dose. Moreover, they need not have an explicit target size, and typically do not fully take into account how much tumor size recently increased or decreased, e.g., they increase the dose by the same amount whether tumor size increased substantially or hugely. We therefore propose a new protocol. As previous protocols, it assumes the existence of a tumor biomarker. Details may be found in Supplementary Material, Section 6, but the basic idea is as follows.

The initial dose is neither zero nor the maximal tolerated dose, but an intermediate dose; e.g., a dose expected to a least stabilize tumor in 75% of patients. The variation of the biomarker is then relatively quickly measured, and a second dose is chosen that differs enough from the first dose to be expected to bring tumor back towards its initial size *N*_0_. From the third measurement on, an educated guess for a stabilizing dose may be made based on simple tumor growth models (see below). The dose delivered until the next measurement is then the estimated stabilizing dose if tumor size is equal to its target, a higher dose if it is above, and a lower dose if it below.

A concrete example is to fix a low and a high threshold, *N*_*l*_ *< N*_0_ and *N*_*h*_ *> N*_0_, positive parameters *γ*_1_ and *γ*_2_, and to deliver the following dose until the next measurement, where *N* is the current estimated tumor size, and *C*_*guess*_ the estimated stabilizing dose:

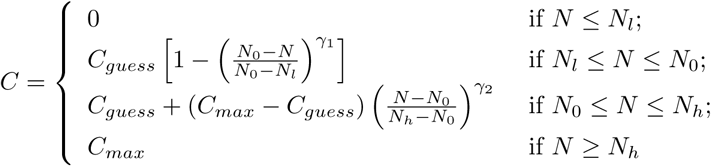

The dose thus varies continuously between 0 and *C*_*max*_ depending on the estimated stabilizing dose and how far tumor size is from target. The parameters *γ*_1_ and *γ*_2_ tune whether the emphasis is on stabilizing tumor size (*γ*_*i*_ *>* 1) or bringing it back to *N*_0_ (*γ*_*i*_ *<* 1). Note that only the ratios *N/N*_0_, *N*_*h*_*/N*_0_ and *N*_*l*_*/N*_0_ matter. This is important when the evolution of the biomarker level in a given patient correlates well with the evolution of tumor size, but the initial level is not a good indicator of initial tumor size.

We conclude by explaining how to find an estimation of the stabilizing dose. Assume that the biomarker levels were measured at time *t*_*k*− 2_, *t*_*k*− 1_ and at the current time *t*_*k*_, leading to estimated tumor doubling times of *DT*_*k*− 2_ between *t*_*k*− 2_ and *t*_*k*− 1_, and *DT*_*k*− 1_ between *t*_*k*− 1_ and *t*_*k*_ (if the tumor is regressing, the doubling time is defined as the opposite of the “halving time”). Denote by *C*_*k*− 2_ and *C*_*k*− 1_ the doses given between *t*_*k*− 2_ and *t*_*k*− 1_, and between *t*_*k*− 1_ and *t*_*k*_, respectively. We show in Supplementary material, Section 6, that a reasonable guess for the stabilizing dose is:

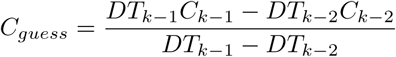

This assumes that the kill-rate is proportional to the dose, as in Model 2. If the kill-rate is assumed proportional to a function *f* (*C*) of the dose, due, e.g., to some saturation effect, then the formula becomes

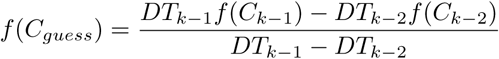

and a guess for the function *f* should also be made. Wrong guesses for this function could lead to systematic under-estimation or over-estimation of the stabilizing dose, but this is compensated by the fact that the protocol takes into account how far tumor size is from target. Moreover, if measurements are frequent enough, learning algorithms may help to determine which models work best to predict tumor evolution in a specific patient, and to gradually improve the estimation of the stabilizing dose. For a more substantial discussion of containment protocols, see Supplementary material, Section 6, and the forthcoming Ph.D. thesis of Jessica Cunningham.

## Discussion

Theoretical support for maximum tolerated dose therapy relies on the assumption that resistant cancer cells are absent [1] or arise only during treatment [2]. Given that many if not most large solid cancers are expected to harbor pre-existing resistance [36], we have sought to build a firm theoretical foundation for understanding when containment strategies are likely to improve on the conventional approach. The logic of containing tumors is fundamentally simple: if some cells are fully resistant to treatment then the only way to fight them is via competition with sensitive cells, where “competition” includes any process that leads to a decrease in the resistant population growth rate due to the presence of sensitive cells. Moreover, given the constraint of maintaining tumor size below a certain threshold, competition is maximized under containment treatment strategies. We have shown that this logic can be formalized and given a rigorous mathematical form in a general setting. It follows that model details are qualitatively irrelevant, provided that resistant cells are highly resistant and that increasing the number of sensitive cells always decreases the resistant population growth rate.

However, identifying conditions under which containment strategies are expected to perform well also emphasizes that the case for containment is weaker when these conditions are not met. For instance, though we checked that taking into account random genetic mutations occurring after treatment initiation would not have substantially affected our results, we did not investigate treatment-induced mutations, nor models where sensitive cells arise from resistant tumor stem cells. We also note that if resistant cells are only partially resistant then switching to MTD before the failure of low dose treatment may be superior to a pure containment strategy. When to switch and whether the difference in outcomes is substantial remains an important topic for further investigation.

In our simple but general framework, the time until tumor size exceeds any particular threshold is maximized by maintaining tumor size precisely at this threshold for as long as there remain sensitive cells. This is true even if resistance has no cellular fitness cost. This suggest that tumor containment experiments and trials should not be restricted to cases where a resistance cost is assumed to exist. Our results also underline a trade-off between maximizing time to progression and maximizing the time at which tumor size becomes higher than some larger threshold. Since clinical evidence supporting containment strategies remains limited, it seems safer to test containing tumors at their initial size, or some relatively low size. If results are convincing, more ambitious strategies aiming at increasing intra-tumor competition by letting the tumor grow to its maximal tolerable size before containing it could be attempted. This maximal tolerable size would not have to be known in advance, but could be discovered during treatment, based on patient’s quality of life.

To implement containment strategies, the nature of the resistance mechanism, the frequency of resistant cells, or other patient specific parameters need not be known, but a tumor burden indicator seems required. In our models, when resistant cells are initially rare, applying a dose close to the initial stabilizing dose throughout typically leads to results similar to containment at the initial size. In practice however, tumor growth is much more irregular. Thus, finding a dose, or schedule, that initially results in tumor stabilization is not enough: regular monitoring and dose adjustment are required. We proposed a new protocol that takes into account how far tumor size is from its target and how much it recently increased or decreased.

Importantly, although the ideal form of containment is impractical, our simulations and theoretical arguments predict that more feasible containment strategies will also improve substantially on maximum tolerated dose (MTD) treatment. These more practical approaches include adaptive therapy [4], which has an important advantage over constant-dose or metronomic protocols, in that the optimal dose need not be known in advance. On the other hand, our theoretical results imply that an on-off implementation of adaptive therapy – as was employed in the only clinical trial of tumor containment to date [12] – may be suboptimal, because it causes tumor size to deviate substantially below the maximum tolerable threshold. Further research is needed to establish optimal dosing protocols in the presence of biological factors not accounted for in our framework, e.g., spatial structure [17].

By deriving explicit formulas for predicted clinical gains due to containment, we have shown that a crucial factor is the intensity of competition between sensitive and resistant cells. For tumors that obey the Gompertzian growth law, clinical gains are predicted to be substantial, at least when resistant cells are initially rare and the initial tumor size is not very small (at least 0.1% of carrying capacity). Less conventional tumor growth models predict either smaller or larger clinical gains. Our findings therefore underscore the need to characterize intratumor cell-cell competition [29]. A useful indicator that could be measured experimentally is the amount by which the resistant population growth rate increases – if at all – upon elimination of sensitive cells.

Although we have investigated various extensions and variants of our basic model, we have not considered all potential clinical costs and benefits of containment. By maintaining a substantial tumor burden, containment might increase risk of metastasis, cancer-induced illness such as cachexia, or emergence of more aggressive tumor clones via mutation [37]. On the other hand, containment has the important advantage of reduced treatment toxicity. Stabilizing tumor size might additionally lead to a more stable tumor microenvironment and better drug delivery, which would be consistent with the finding that, in preclinical trials in mice, tumor size could be stabilized using progressively lower doses [11]. Further experimental and theoretical research is needed to clarify whether the benefit of containment in terms of prolonging survival always outweighs its potential downsides. Notwithstanding these important caveats, our findings generally strengthen the case for conducting further experimental and clinical trials of tumor containment strategies.

## Acknowledgments

We are grateful to Pierre Lissy, Sébastien Benzekry, and especially Joel Brown for very helpful discussions, and to Philippe Ear for preliminary versions of some figures. This research benefited from the support of the FMJH “Program Gaspard Monge for optimization and operation research and their interactions with data science” and from the support from EDF, Thales, Orange and Criteo. The project was initiated at the Lorentz Center Workshops “Game Theory and Evolutionary Biology” and “Understanding Cancer Through Evolutionary Game Theory”.

## Notes

### Competing Interest Statement

The authors have declared no competing interest.

### Summary of Updates

Additional and revised figures; methods more fully described in the main text; some new results.

