## Supplementary material for "The logic of containing tumors"

### THE LOGIC OF CONTAINING TUMORS: SUPPLEMENTARY MATERIAL

#### Contents

1. Partial survey of related works 2
2. Qualitative comparison of treatments under a general model 3
2.1. Model, treatments, notation 3
2.2. Informal description of results 5
2.3. Formal results 5
2.4. The case $N_0 > N_{tol}$ 7
2.5. Reminder on differential equations and proof of Proposition 1 8
3. Explicit formulas 9
3.1. The purely density-dependent case 10
3.2. Frequency-dependent models and models with resistance costs 12
3.3. Application to Gompertzian growth 13
4. Comparison between treatments 15
4.1. Idealized treatments 15
4.2. Containment and ideal containment 17
4.3. MTD and ideal MTD 18
4.4. MTD and containment 19
4.5. Impact of varying $C_{max}$ 19
4.6. Constant dose and containment 21
5. Impact of resistance costs on the best possible outcome and on clinical benefits of containment 22
5.1. Justification of main text Figure 4b 22
5.2. Approximate formula for the benefit of containment with resistance costs 22
6. Containment at the initial size in practice 24
6.1. Some simple protocols 24
6.2. A new protocol 25
6.3. How to make an educated guess for the current stabilizing dose? 27
7. Taking mutations into account 28
7.1. Variant 1: Late first mutation. 29
7.2. Variant 2: Birth-death model 29
7.3. Variant 3: Slowly growing resistant cells 30
7.4. Discussion: links with Hansen et al. (2017) 31
References 32

This supplementary material is organized as follows. Section 1 recalls a number of previous models related to our work. Section 2 studies Model 1 from the main text. For this general model, we qualitatively compare reference treatments and an arbitrary treatment, in terms of sensitive and resistant population sizes and the time until tumor size exceeds an arbitrary threshold. This time is shown to be maximized by an idealized version of containment at this threshold. Different choices of threshold lead to results on time to progression, time to treatment failure, or survival time. The proofs are based on variants of Gronwall's inequalities.

Section 3 considers various density- and frequency-dependent models. Explicit formulas are derived for the time at which tumor size exceeds an arbitrary threshold under various treatments. This result is then applied to Gompertzian growth (Model 2 in the main text). Building on these findings, Section 4 compares reference treatments qualitatively and quantitatively. Section 5 studies the impact of resistant costs on the best possible outcome, and on the clinical benefits of containment. This section provides an approximate formula for time to treatment failure under containment or ideal containment in the presence of resistance costs. Section 6 discusses possible protocols to implement containment at a target size. Finally, Section 7 shows that accounting for mutations from sensitive to resistant cells that occur after treatment initiation does not affect substantially the time to progression for the containment treatment (nor time to treatment failure, nor survival time). This justifies our neglecting such mutations in the bulk of our analysis.

#### 1. Partial survey of related works

We recall here some of the models most related to our work.

*Lotka-Volterra and density-dependent models.* Zhang *et al.* (2017) [1] consider a Lotka-Volterra model with three types of tumor cells (two sensitive and one resistant to treatment), and carrying capacities that depend on whether treatment is on or off. Through simulations, they compare an intermittent containment treatment (maintaining the tumor between its initial size $N_0$ and $N_0/2$) to no treatment, maximal tolerated dose (MTD) and a form of metronomic therapy with an induction period. Cunningham *et al.* (2018) [2] extend this model to allow for intermediate dose treatments. This extended model could be simplified by grouping the two types of sensitive cells, leading to a two-type Lotka-Volterra model of the form:

$$\dot{s} = \rho_s s \left(1 - \frac{(\alpha_1 s + \alpha_2 r)}{K_s}\right) \tag{1}$$

$$\dot{r} = \rho_r r \left(1 - \frac{(r + \beta s)}{K_r}\right) \tag{2}$$

with $K_r$ independent of treatment, and $K_s$ linearly varying as a function of the dose between $K_{max} = K_r$ and a much lower value $K_{min} = K_r/100$. The competition coefficient $\beta$ is assumed less than 1, so the tumor cannot be stabilized at a size smaller than $K_r$.

Carrère (2017) [3] studies a similar two-type Lotka-Volterra model but with a higher impact of sensitive cells on resistant cells, and a different treatment-induced death term. She discusses how to optimally control the tumor for various objective functions. Carrère and Zidani (2019) [4] consider an extension of Carrère's model, in particular adding uncertainties on some parameters, and use optimal control to study how to bring and maintain tumor size below a certain threshold. Pouchol *et al.* (2018) [5] uses optimal control techniques to study a model with infinitely many types and two types of drug: cytostatic and cytotoxic.

A precursor of such optimal control approaches for models with intratumor competition is Martin *et al.* (1992) [6]. They consider, with some approximations, a general density-dependent two-type model with mutations, and study how to optimally tune tumor size in order to maximize survival time. Applications are made to exponential, logistic, and Gompertzian growth. The analysis is extended to combination chemotherapies in [7]. Building on this seminal work, Hansen *et al.* (2017) [8] focus on the logistic and Gompertzian case, and compare two treatments: a form of ideal containment, and elimination of sensitive cells, in a context broader than cancer. We borrowed the term containment from them.[1] In a follow-up paper, Hansen *et al.* (2020) [9] compare experimentally and theoretically the growth of a partially resistant strain of *E. coli*, with or without adding competing sensitive bacteria. Hansen and Read (2020) [10] consider a stochastic birth-death model allowing for a positive probability of cure even when resistant cells are initially present. They discuss the trade-off between a higher probability of cure and a shorter time to progression in case of failed cure (see also [11]).

Monro and Gaffney (2009) [11], in a paper published a few months before the first article on adaptive therapy (Gatenby *et al.* 2009 [12]), study constant dose treatments in a two-type model with Gompertzian growth and no resistance cost. This is the model we use for simulations (except that we neglect mutations). They show that reducing the dose or delaying treatment may increase survival time.[2]

*Frequency-dependent models.* Another line of models assumes frequency-dependent competition. For instance, Silva *et al.* (2012) [13] considers a discrete-time, difference equation model, which in a continuous-time, differential equation framework would take the form

$$\begin{aligned} \dot{s}/s &= \rho_s \frac{s}{s+r} - \lambda_s C \\ \dot{r}/r &= \rho_r \frac{r}{s+r} - \lambda_r C \end{aligned}$$

[1] Containment in Hansen *et al.* (2017) [8] is as our ideal containment treatment except that they also allow tumor size to *increase* instantly from its initial size to a possibly higher size, at which the tumor is then maintained. We only allow for an instantaneous *decrease* of tumor size.

[2] Gatenby *et al.* (2009) [12] also contains a model, but substantially different and more involved that the models we use, and we will not discuss it here.

where $C$ is a measure of treatment intensity, $\lambda_s$ and $\lambda_r$ represent sensitivity to treatment of sensitive and resistant cells, and the growth-rate parameters $\rho_r$ and $\rho_s$ vary as a function of some auxiliary treatment. In this model, as the frequency of resistant cells $x_r = \frac{r}{s+r}$ approaches zero, the relative fitness of resistant cells approaches zero, allowing for a huge, unbounded advantage of containment over MTD. Bacevic *et al.* (2017) [14] considers a similar frequency-dependent model, but where the resistant population growth rate is proportional to a function $f(x_r)$ with $f$ bounded away from zero. The relative gain of containment compared to MTD (that is, the ratio of the time it takes under these treatments for the tumor to reach a given size) is then bounded by $1/f(0)$. Bacevic *et al.* also studies models mixing frequency dependence and Gompertzian, density-dependent growth, and models where the carrying capacity is dynamic and reduced by treatment, in the spirit of Hahnfeldt *et al.* (1999) [15]. In the latter case, resistant cells are, indirectly, still partially sensitive to treatment, and our analysis does not apply.

Some features of these models are summed up in Supplementary Table 1. A number of other interesting approaches are less connected to our work. For instance, Gallaher *et al.* (2018) [16] studies a spatial dose modulation model through simulations. West *et al.* (2018) [17] considers game theoretical models with sensitive and resistant cells, but also normal cells as a third type. Ledzewicz and Schättler (2019) [18] review optimal control models for heterogeneous tumors with more or less resistant cells, but the objective functions are different and the models studied do not take into account competition between cell types. This mini-review is far from exhaustive and we apologize for all the fine works that are not mentioned above.

The main differences between our work and the bulk of the adaptive therapy literature are the generality of our results, and in particular the fact that we provide analytical results that apply to many different models instead of relying on simulations of a particular model. The main differences with the bulk of the optimal control literature (with some exceptions, e.g., Martin *et al.* (1992) [6, 7], Carrère and Zidani (2019) [4]) are the focus on intra-tumor competition, our objective function (maximizing the time at which tumor exceeds a given size), and the fact that our proofs rely on basic variants of Gronwall's lemma rather than the heavier optimal control machinery.

#### 2. Qualitative comparison of treatments under a general model

2.1. **Model, treatments, notation.** We study here Model 1 from main text. We first recall it, and clarify assumptions that are only informally described in the main text. The model reads

$$\begin{cases} \dot{S}(t) = S(t)g_s(S(t), R(t), C(t)) & ; \quad S(0) = S_0 \geq 0 \\ \dot{R}(t) = R(t)g_r(S(t), R(t)) & ; \quad R(0) = R_0 > 0 \end{cases} \tag{3}$$

where $S(t)$, $R(t)$ are the number of sensitive and fully resistant cells, respectively, and $C(t)$ is the drug dose or more generally the treatment level at time $t$.[3] The total tumor population size is $N(t) = S(t) + R(t)$, with initial value $N_0 = S_0 + R_0$.

*Model assumptions.* The key assumptions are that resistant cells are fully resistant and that $g_r$ is strictly decreasing in $S$. We also assume that $g_s$ is non-increasing in $R$ and strictly decreasing in $C$, that as long as the patient is alive ($N < N_{crit}$), the size of an untreated or fully resistant tumor strictly increases, and that, for any treatment level $C$, the function

$$R \to (N_{tol} - R)g_s(N_{tol} - R, R, C) + Rg_r(N - R, R)$$

is increasing on $[0, N_{tol}]$. This last assumption ensures that if a tumor treated at MTD increases beyond the maximum tolerable level $N_{tol}$ then it will never become smaller again. It also implies that the treatment level required to stabilize a tumor of size $N_{tol}$ increases with the frequency of resistant cells.

Finally, we make technical assumptions: functions $g_r$ and $g_s$ are continuously differentiable on the relevant domain ($N > 0$, $C \geq 0$); solutions are defined for all $t \geq 0$; and function $C$ is continuous on $[0, +\infty)$ (this simplifies proofs but our main results also hold for piecewise continuous treatments, and discontinuities in the sensitive population size).

*Objective.* Treatment is said to fail when tumor size exceeds a *maximum tolerable size* $N_{tol}$. This size need not be known in advance but could instead be identified during treatment. Our treatment objective is then to maximize *time to treatment failure*: the largest time $\tau$ such that $N(t) \leq N_{tol}$ on $[0, \tau]$. To simplify

[3] The difference between drug dose and treatment level is two-fold: first, the model applies to treatments such as radiotherapy that are not naturally described as drugs; second, we neglect pharmacodynamics and pharmacokinetics. As already pointed out by Norton and Simon (1977) [19], "depending on the type of therapy used and such factors as route of administration or concurrent medication, [treatment level] may be related to the dose administered in a complicated fashion".

Supplementary table 1. **Features of adaptive therapy and containment models.** In the treatment column, "containment" refers to various implementations of the general containment idea.

| **Study** | **Types** | **Treatment** | **Competition type** | **Methods** | **Other features** |
|---|---|---|---|---|---|
| Martin *et al.* (1992) [6] | 2 | any | density-dependent; Gompertzian; Lotka-Volterra | optimal control | no resistance cost; mutations |
| Monro & Gaffney (2009) [11] | 2 | constant dose; delaying treatment | Gompertzian | simulations | no resistance cost; mutations |
| Gatenby *et al.* (2009) [12] | various | containment; metronomic; MTD | unconventional | analytical results; simulations | resistance cost; microenvironmental feedback |
| Silva *et al.* (2012) [13] | 2 | containment; MTD | specific frequency-dependent | simulations | bolus doses; manipulation of resistance cost |
| Hansen *et al.* (2017) [8] | 2 | containment; ideal MTD | Lotka-Volterra; Gompertzian | analytical results | resistance cost or not; mutations |
| Carrère (2017) [3] | 2 | any | Lotka-Volterra | optimal control | resistance cost |
| Bacevic *et al.* (2017) [14] | 2 | containment; MTD | specific frequency & density-dependent | simulations; analytical results | bolus doses; resistance cost; partial resistance |
| Zhang *et al.* (2017) [1] | 3 | containment; metronomic; MTD | Lotka-Volterra | simulations | clinical data |
| Pouchol *et al.* (2018) [5] | any | any | Lotka-Volterra | optimal control | cytostatic and cytotoxic drugs |
| Cunningham *et al.* (2018) [2] | 3 | any | Lotka-Volterra | numerical optimal control | fixed total dose |
| Hansen *et al.* (2020) [9] | 2 | containment; ideal MTD | Lotka-Volterra | simulations | corresponding *in vitro* experiments |
| Hansen and Read (2020) [10] | 2 | containment; ideal MTD | Lotka-Volterra | simulations; analytical results | possibility of cure |
| Carrère and Zidani (2019) [4] | 2 | any | Lotka-Volterra | optimal control | tolerable tumor size |
| Current study | 2 | any | general frequency- & density-dependent | simulations; analytical results; comparison principle | resistance cost or not; tolerable tumor size |

the exposition, we assume by default that the maximum tolerable size is no smaller than the initial size: $N_{tol} \geq N_0$ (the case $N_{tol} < N_0$ is discussed in Section 2.4). We make no other assumption about $N_{tol}$. In particular, taking $N_{tol} = N_0$ leads to results on *time to progression*, and setting $N_{tol}$ to be the lethal tumor burden gives results on *survival time*.

*Treatments.* We consider the following treatments (with corresponding subscript in parenthesis):

- No treatment (noTreat): $C(t) = 0$ throughout.
- MTD (MTD): $C(t) = C_{max}$ throughout.
- delayed MTD (del-MTD): does not treat until $N = N_{tol}$, then $C(t) = C_{max}$ for ever.
- containment at $N_{tol}$ (Cont): does not treat until $N = N_{tol}$, then stabilizes tumor size at $N_{tol}$ as long as possible with a dose $C(t) \leq C_{max}$, then treats at $C_{max}$ (Figs. 1d, 1e).
- intermittent containment between $N_{tol}$ and $N_{min} < N_{tol}$ (Int): does not treat until $N = N_{tol}$, then treats at $C_{max}$ until $N = N_{min}$, and iterates as long as possible, as in [1] (Fig. 1g).[4]

We also consider idealized versions, which may be thought of as relaxing the constraint $C(t) \leq C_{max}$:

- ideal MTD (idMTD): instantly eliminates sensitive cells ($S(t) = 0$ for all $t > 0$.)[5]
- delayed ideal MTD (del-idMTD): does not treat until $N = N_{tol}$, then instantly eliminates sensitive cells.
- ideal containment at $N_{tol}$ (idCont): does not treat until $N = N_{tol}$, then stabilizes tumor size at $N_{tol}$ as long as some sensitive cells remain.
- ideal intermittent containment (idInt): as intermittent containment, except that upon reaching $N_{tol}$, tumor size is instantly reduced to $N_{min}$ (or to $R$, if $R > N_{min}$).

[4]Our results actually hold for any other way of maintaining tumor size between $N_{min}$ and $N_{max}$.

[5]This treatment is called "aggressive treatment" or "elimination of sensitive cells" by Hansen *et al.* (2017, 2020) [8, 9]. We may think of this as a hypothetical infinite-dose treatment, or more precisely a treatment with an infinite kill-rate. Alternatively, in an experimental setting, ideal MTD may correspond to the initial condition of a fully resistant population of cells (cf. Hansen *et al.*, 2020 [9]).

We compare these treatments between themselves and to an arbitrary alternative treatment, that we only assume regular enough to avoid technical issues. To simplify some statements, all treatments are assumed to treat at $C_{max}$ after treatment failure.

*Notation.* Times to treatment failure are denoted by $t_{noTreat}$, $t_{MTD}$, $t_{del-MTD}$, $t_{Cont}$ and $t_{Int}$, respectively, for non-idealized treatments; $t_{idMTD}$, $t_{del-idMTD}$, $t_{idCont}$ and $t_{idInt}$, for idealized treatments; and $t_{alt}$ for the alternative treatment. Similar subscripts are used to refer to treatment level, and to the sensitive, resistant, and total tumor sizes under these treatments (e.g., $C_{alt}(t)$, $S_{alt}(t)$, $R_{alt}(t)$, and $N_{alt}(t)$ for the alternative treatment).

2.2. **Informal description of results.** Intuitively, if resistant cells are fully resistant, then the only way to fight them is via competition with sensitive cells. Our aim is to turn this intuition into rigorous mathematical results allowing to compare the effect of various treatments in Model 1. The key result (Proposition 1) is that, in our model, more sensitive cells lead to fewer resistant cells, and this implication can be formally proven. Similarly, a larger tumor burden – or a lower dose – leads to fewer resistant cells (and more sensitive cells). It follows that eliminating sensitive cells maximizes the resistant population, and minimizes time to treatment failure among treatments that fully eliminate sensitive cells before failing (Proposition 2). Conversely, by maintaining tumor size as high as possible before failing, ideal containment maximises time to treatment failure (Proposition 3).

Under the constraint $C(t) \leq C_{max}$, containment does not exactly maximize time to treatment failure, because switching to MTD shortly before treatment failure would result in a small delay. However, any treatment that switches to MTD after failing would lead to a larger resistant population at all times, hence typically a larger long-term tumor burden (Proposition 4). In particular, though containment fails before its idealized version, the constraint $C(t) \leq C_{max}$ leads to a lower resistant population than in ideal containment.

A yet more realistic protocol is intermittent containment, which aims at maintaining tumor burden between two thresholds [1]. Consistent with intuition, intermittent containment is intermediate between containment at the lower threshold and containment at the higher threshold in terms of sizes of resistant and sensitive populations, and in terms of time to treatment failure in the idealized case (Propositions 5 and 6). Similarly, delaying treatment before treating at MTD is intermediate between MTD and intermittent containment (Proposition 7, which also sums up the comparison between all reference treatments).

Finally, Section 2.4 considers the case in which the initial tumor burden is above the tolerable threshold ($N_0 > N_{tol}$). The ideal containment strategy then first reduces tumor size to the maximum tolerable size (if $R_0 < N_{tol}$), and then stabilizes the size of the tumor as long as it is not fully resistant. This strategy is shown to maximize the time for which tumor size is no larger than the maximal tolerable size (Proposition 8).

2.3. **Formal results.** We begin with a key result, proved in Section 2.5. It shows that keeping more sensitive cells, a larger tumor burden, or treating less, leads to fewer resistant cells.

**Proposition 1.** *(key result) Let $0 \leq t_0 \leq t_1$. Consider two continuous treatment level functions $C_1$, $C_2$, with associated tumor subpopulation sizes $(S_1, R_1)$ and $(S_2, R_2)$, satisfying* (3)*. Let $N_i = S_i + R_i$, $i = 1, 2$, denote total tumor size. Assume that at time $t_0$, the resistant population is non-larger, and the sensitive population no-smaller under the first treatment than under the second: i) $R_1(t_0) \leq R_2(t_0)$; and ii) $S_1(t_0) \geq S_2(t_0)$. Assume moreover that between $t_0$ and $t_1$, at least one of the following conditions holds: under treatment 1,*

*iiia) the sensitive population is larger: $\forall t \in [t_0, t_1], S_1(t) \geq S_2(t)$;*
*or iiib) total tumor size is larger: $\forall t \in [t_0, t_1], N_1(t) \geq N_2(t)$;*
*or iiic) treatment level is lower: $\forall t \in [t_0, t_1], C_1(t) \leq C_2(t)$.*
*Then, for all $t$ in $[t_0, t_1]$, $R_1(t) \leq R_2(t)$ and $S_1(t) \geq S_2(t)$.*

It follows that ideal MTD (or MTD, under the constraint $C(t) \leq C_{max}$) maximizes the resistant population size (see also [7], [8]). Ideal MTD also minimizes time to treatment failure among all treatments that eliminate sensitive cells before failing.[6]

**Proposition 2.** *(comparison with no treatment, MTD and ideal MTD)*

[6]The latter result should be seen as a comparison between idealized treatments. Indeed, under the constraint $C(t) \leq C_{max}$, the sensitive population size would never be exactly 0.

a) *For all times* $t \geq 0$, $R_{noTreat}(t) \leq R_{alt}(t) \leq R_{MTD}(t) \leq R_{idMTD}(t)$ *and* $S_{idMTD}(t) \leq S_{MTD}(t) \leq S_{alt}(t) \leq S_{noTreat}(t)$, *where the comparisons between MTD and the alternative treatment are only valid if* $C_{alt}(t) \leq C_{max}$ *for all* $t \geq 0$.

b) *If* $S_{alt}(t_{alt}) = 0$, *then* $t_{alt} \geq t_{idMTD}$

*Proof.* a) Immediate by Proposition 1; b) if $S(t_{alt}) = 0$, then $N_{tol} = N_{alt}(t_{alt}) = R_{alt}(t_{alt}) \leq R_{idMTD}(t_{alt})$ by a), hence $t_{idMTD} \leq t_{alt}$. □

Proposition 1 also implies that ideal containment leads to a smaller resistant population than under any alternative treatment that has not yet failed, and maximizes time to treatment failure.

**Proposition 3.** *(comparison with ideal containment)*

a) *For all* $t$ *in* $[0, t_{alt}]$, $R_{idCont}(t) \leq R_{alt}(t)$ *and* $S_{idCont}(t) \geq S_{alt}(t)$.

b) $t_{idCont} \geq t_{alt}$

*Proof.* a) Let $t_0$ be the first time such that $N_{idCont}(t_0) = N_{tol}$. On $[0, t_0]$, ideal containment does not treat, so these inequalities hold by Proposition 2. If $t_0 \leq t_{alt}$, then on $[t_0, t_{alt}]$, $N_{idCont}(t) \geq N_{alt}(t)$. Since the inequalities hold for $t = t_0$, it follows from Proposition 1 that they still hold on $[t_0, t_{alt}]$.

b) Therefore, $N_{tol} = N_{alt}(t_{alt}) \geq R_{alt}(t_{alt}) \geq R_{idCont}(t_{alt})$, which implies that $t_{idCont} \geq t_{alt}$, since under ideal containment, failure occurs when $R = N_{tol}$. □

Under a maximal instantaneous dose constraint, containment at $N_{tol}$ does not exactly maximize time to treatment failure. Indeed, contrary to what happens with ideal containment, there are still sensitive cells at treatment failure. For this reason, switching to MTD slightly before containment fails would slightly delay treatment failure.[7] However, containment leads to a lower resistant population than any treatment that treats at $C_{max}$ after failing, in particular than ideal containment.

**Proposition 4.** *(containment) For all* $t \geq 0$, $R_{Cont}(t) \leq R_{alt}(t)$ *and* $S_{Cont}(t) \geq S_{alt}(t)$.

*Proof.* For $t \leq t_{alt}$, the proof is as for ideal containment. Moreover, by assumption, for all $t \geq t_{alt}$, $C_{alt}(t) = C_{max} \geq C_{Cont}(t)$. Thus by Proposition 1, the desired inequalities still hold at all later times. □

We now compare intermittent containment between $N_{tol}$ and $N_{min} < N_{tol}$ to containment at the higher threshold $N_{tol}$ and containment at the lower threshold $N_{min}$. The latter let tumor grow to $N_{min}$ (or treats at $C_{max}$ until $N = N_{min}$ if $N_0 > N_{min}$), then stabilizes tumor size at $N_{min}$ as long as possible with a dose $C(t) \leq C_{max}$, and then treats at $C_{max}$. In the idealized version, tumor size is stabilized at $N_{min}$ as long as some sensitive cells remain (and tumor size is instantly reduced to $N_{min}$ at time 0 if $N_0 > N_{min}$). The subscripts used for containment and ideal containment at $N_{min}$ are ContNmin and idContNmin, respectively.

The next result shows that idealized intermittent containment is, in a precise sense, intermediate between ideal containment at the lower and at the higher level.

**Proposition 5.** *(intermittent versus continuous containment: idealized treatments)*

a) *For all* $t \geq 0$, $R_{idCont}(t) \leq R_{idInt}(t) \leq R_{idContNmin}(t)$ *and* $S_{idContNmin}(t) \leq S_{idInt}(t) \leq S_{idCont}(t)$

b) $t_{idContNmin} \leq t_{idInt} \leq t_{idCont}$.

c) *For all* $t \geq t_{idInt}$, $N_{idCont}(t) \leq N_{idInt}(t) \leq N_{idContNmin}(t)$.

*Proof.* The comparison with ideal containment at $N_{tol}$ (idCont) follows from Proposition 3. Let us compare ideal intermittent containment and ideal containment at $N_{min}$. Let

$$t^* = \sup\{\tau \geq 0, N_{idContNmin}(t) \leq N_{idInt}(t) \text{ on } [0, \tau]\}$$

and

$$\hat{t} = \sup\{t \geq 0, N_{idContNmin}(t) \leq N_{min}\}$$

and note that $\hat{t} \leq t^*$. For $t \in [0, t^*]$, it follows from Proposition 1 that $R_{idContNmin}(t) \geq R_{idInt}(t)$ and $S_{idContNmin}(t) \leq S_{idInt}(t)$. In particular, this holds at $\hat{t}$. But for $t \geq \hat{t}$, $S_{idContNmin}(t) = 0 \leq S_{idInt}(t)$. Therefore, the above inequalities are still valid at all later times by Proposition 1, and are thus valid for all

[7] The omitted proof of this result is easy. It simply exploits the fact that, in the short run, increasing treatment decreases both the sensitive population and tumor size. In the long run, this would lead to a quicker development of resistant cells and typically a larger tumor size, but increasing treatment only shortly before $t_{Cont}$ ensures that the short-run effect dominates until $t_{Cont}$.

positive times. In particular, at $t_{idInt}$, $N_{tol} = R_{idInt} \leq R_{idContNmin}$, hence $t_{idContNmin} \leq t_{idInt}$. Finally, for $t \geq t_{idInt}$, $N_{idInt} = R_{idInt} \leq R_{idContNmin} \leq N_{idContNmin}$. This concludes the proof. □

The next proposition is a partial analog for non-idealized treatments. The proof uses that containment at $N_{min}$ treats at $C_{max}$ when $N > N_{min}$.

**Proposition 6.** *(intermittent containment: bounded instantaneous dose)* *For all* $t \geq 0$, $R_{Cont}(t) \leq R_{Int}(t) \leq R_{ContNmin}(t)$ *and* $S_{ContNmin}(t) \leq S_{Int}(t) \leq S_{Cont}(t)$

*Proof.* The comparison with containment at $N_{tol}$ follows from Proposition 4. If $N_{min} \geq N_0$, then the comparison with containment at $N_{min}$ is similar to the comparison with ideal containment at $N_{min}$ in Proposition 5, but replacing $S_{idContNmin} \leq S_{idInt}$ by $C_{ContNmin} \geq C_{Int}$ once $N_{ContNmin} > N_{min}$. If $N_{min} < N_0$, then as long as $N_{Contmin} > N_0$, $C_{Contmin} = C_{max} \geq C_{Int}$, hence the result holds by Proposition 1; if at some time $\tilde{t}$, $N_{Contmin} = N_{min}$, the proof that the required inequalities hold also for $t \geq \tilde{t}$ is as in the case $N_0 \geq N_{min}$. □

Our final result compares all reference treatments, including delayed MTD and its idealized version.

**Proposition 7.** *(comparison between all reference treatments) For all* $t \geq 0$*:*

a) $S_{idMTD}(t) \leq S_{del-idMTD}(t) \leq S_{idInt}(t) \leq S_{idCont}(t) \leq S_{noTreat}(t)$
b) $R_{idMTD}(t) \geq R_{del-idMTD}(t) \geq R_{idInt}(t) \geq R_{idCont}(t) \geq R_{noTreat}(t)$
c) $t_{idMTD} \leq t_{del-idMTD} \leq t_{idInt} \leq t_{idCont}$
d) $S_{MTD}(t) \leq S_{del-MTD}(t) \leq S_{Int}(t) \leq S_{Cont}(t) \leq S_{noTreat}(t)$
e) $R_{MTD}(t) \geq R_{del-MTD}(t) \geq R_{Int}(t) \geq R_{Cont}(t) \geq R_{noTreat}(t)$

*Moreover, for all* $t \geq t_{idCont}$, $N_{idMTD}(t) \geq N_{del-idMTD}(t) \geq N_{idInt}(t) \geq N_{idCont}(t)$

*Proof.* a) The first two inequalities are immediate from the definition of these treatments. The inequality $S_{idInt}(t) \leq S_{idCont}(t)$ was proved in Proposition 5. The last inequality is immediate from Proposition 1.

b) Immediate from a) and Proposition 1.

c) Immediate from b), since for idealized treatments, $S = 0$ when treatment fails.

d) The first two inequalities are immediate from the definition of these treatments and Proposition 1 (since $C_{MTD}(t) \geq C_{del-idMTD}(t) \geq C_{Int}(t)$ for all $t$). The inequality $S_{Int}(t) \leq S_{Cont}(t)$ was proved in Proposition 6. The last inequality is immediate from Proposition 1.

e) Immediate from d) and Proposition 1.

Finally, the last result is due to b) and to the fact that, by c) and definition of reference idealized treatments, for $t \geq t_{idCont}$, $S = 0$ hence $N = R$ for all these treatments. □

2.4. **The case $N_0 > N_{tol}$.** It could be that at the beginning of treatment, tumor size is already intolerable, that is, $N_0 > N_{tol}$. In that case, maximizing the time at which treatment fails is not an appropriate objective, since, with our definition of treatment failure, treatment fails before beginning. Another possible objective is to maximize the total time spent at tumor sizes below $N_{tol}$; that is, the quantity

$$\tau = \int_0^{+\infty} \mathbb{1}_{N(t) \leq N_{tol}} \, dt$$

where $\mathbb{1}_{N(t) \leq N_{tol}} = 1$ if $N(t) \leq N_{tol}$ and 0 otherwise.

Containment could be thought of as first treating at MTD until tumor size is tolerable, and then trying to stabilize tumor size at $N_{tol}$ for as long as possible. In the idealized version – our definition of ideal containment when $N_0 > N_{tol}$ – tumor size is instantly reduced from $N_0$ to $N_{tol}$, and then stabilized at this size as long as $R(t) \leq N_{tol}$. Our next result is that ideal containment is optimal for the above objective, under the additional assumption that the resistant population may be slowed down by the presence of sensitive cells, but nonetheless keeps increasing. This is equivalent to assuming that any tumor containing fully resistant cells is eventually lethal.

The intuition is as follows: first, since the resistant population keeps growing, tumor size should be reduced as quickly as possible, to minimize the size of the resistant population when tumor size becomes tolerable. For the same reason, once the tumor burden is tolerable, there is no advantage in letting tumor burden become temporarily intolerable (this would not contribute to the time spent with a tolerable tumor burden, and while $N > N_{tol}$, the resistant population would still grow, making the situation worse when going back to $N \leq N_{tol}$). It follows that, to be optimal, it suffices to bring tumor size back to $N_{tol}$ as quickly as possible, and then maximize time to treatment failure from that point on. This is precisely what ideal containment does.

**Proposition 8.** *Let $\tau_{idCont}$ and $\tau_{alt}$ denote the total time spent at tumor sizes below $N_{tol}$ under ideal containment and an alternative treatment, respectively. Assume that as long as the patient is alive, the resistant population keeps growing: for all $R \leq N \leq N_{crit}$, $g_r(R, N-R) > 0$, where $N_{crit}$ is the lethal tumor size. Then $\tau_{idCont} \geq \tau_{alt}$.*

*Proof.* First note that, since $R'_{idCont} = g_r(R_{idCont}, N_{tol} - R_{idCont})$ for $0 < t \leq \tau_{idCont}$:

$$\tau_{idCont} = \int_0^{\tau_{idCont}} 1\, dt = \int_0^1 \frac{R'_{idCont}(t)}{g_r(R_{idCont}(t), N_{tol} - R_{idCont}(t))}\, dt = \int_{R_0}^{N_{tol}} \frac{du}{g_r(u, N_{tol}-u)} \tag{4}$$

where we made the change of variables $u = R_{idCont}(t)$ and used that $R_{idCont}(0) = R_0$ and $R_{idCont}(\tau_{idCont}) = N_{tol}$. Second, let $t_0$ and $t_1$ be the first and last times such that $N_{alt}(t) = N_{tol}$ under the alternative treatment. For all $t \leq t_1$, $R_{alt}(t) \leq N_{tol}$ and

$$\mathbb{1}_{N_{alt}(t)\leq N_{tol}} = \frac{R'_{alt}(t)}{g_r(R_{alt}(t), N_{alt}(t) - R_{alt}(t))}\mathbb{1}_{N_{alt}(t)\leq N_{tol}} \leq \frac{R'_{alt}(t)}{g_r(R_{alt}(t), N_{tol} - R_{alt}(t))}$$

Indeed, the Left-Hand-Side is 0 if $N_{alt}(t) > N_{tol}$, and is not larger than the Right-Hand-Side otherwise, since $R'_{alt}(t)$ is positive, and $g_r$ is positive and decreasing in its second argument. Therefore, the change of variable $u = R_{alt}(t)$ leads to:

$$\tau_{alt} = \int_{t_0}^{t_1} \mathbb{1}_{N_{alt}(t)\leq N_{tol}}\, dt \leq \int_{t_0}^{t_1} \frac{R'_{alt}(t)}{g_r(R_{alt}(t), N_{tol} - R_{alt}(t))} dt = \int_{R(t_0)}^{R(t_1)} \frac{du}{g_r(u, N_{tol}-u)} \leq \tau_{idCont}$$

by (4), since $R(t_0) \geq R_0$ and $R(t_1) \leq N_{tol}$. □

A similar reasoning could also be applied to the case $R_0 < N_{tol}$. The result is then that ideal containment not only maximises the first time at which tumor burden becomes larger than $N_{tol}$, but also maximizes the total time spent at tumor sizes below $N_{tol}$ (that is, even considering treatments that would temporarily let tumor grow above $N_{tol}$, then reduce tumor size below this threshold, any number of times, ideal containment maximizes the total time spent at tumor sizes non-larger than $N_{tol}$). This requires the additional assumption that the resistant population may be slowed down by sensitive cells, but keeps increasing as long as the patient is alive, even for very large sensitive population sizes. Otherwise, a possible strategy would be to first let the sensitive population grow so much that the size of the resistant population decreases, and start treating heavily only when the resistant population has almost been eliminated. If this allows obtaining a tumor of size $N_{tol}$ with a resistant population smaller than $R_0$, then this allows obtaining a larger total time spent at tumor sizes below $N_{tol}$. Similar strategies are discussed by Carrère (2017), Pouchol *et al.* (2018), and Carrère and Zidani (2019) [3–5].

##### 2.5. Reminder on differential equations and proof of Proposition 1.

For completeness, we recall differential equation tools used to prove Proposition 1, which can be found in any good advanced textbook. The reader familiar with differential equations and Gronwall's inequalities (which we call here "comparison principles") can jump to Section 2.5.2.

2.5.1. *Reminder on differential equations.* Consider the differential equation

$$\dot{x}(t) = f(t, x(t)) \tag{5}$$

with $f : \mathbb{R}^2 \to \mathbb{R}$. Assume that:

- $f$ is continuous, and admits a continuous partial derivative $\partial f/\partial x$ with respect to its second variable.
- there exist constants $A$ and $B$ such that $|f(t,x)| \leq A|x| + B$ for all $(t,x)$ in $\mathbb{R}^2$.

This ensures that for any $(t_0, x_0)$ in $\mathbb{R}^2$, there is a unique solution such that $x(t_0) = x_0$, and that this solution is defined for all times. The first part also implies that a solution starting below another stays below it.

**Property 9.** *Let $x$ and $y$ be solutions of* (5). *If there exists a time $t_0$ such that $x(t_0) < y(t_0)$, then $x(t) < y(t)$ for all $t$ in $\mathbb{R}$.*

A subsolution of (5) is a differentiable function $u$ such that $u'(t) \leq f(t, u(t))$. A supersolution is a differentiable function $u$ such that $u'(t) \geq f(t, u(t))$. A solution is both a subsolution and a supersolution. The most important tool for our proofs is the following *comparison principle*, a variant of Gronwall's lemma. It says that if a subsolution starts below a solution (or a supersolution), it stays below at all later times (strictly so if it starts strictly below).

**Property 10.** (comparison principle) *Let $t_0 \in \mathbb{R}$. Let $u$ be a subsolution and $v$ a supersolution of* (5), *defined at $t_0$. Assume that $u(t_0) \leq y(t_0)$. Then for all $t \geq t_0$ such that both $u$ and $v$ are defined, $u(t) \leq y(t)$, with a strict inequality if $u(t_0) < v(t_0)$.*

Finally, let $\phi_t(x_0)$ denote the value $x(t)$ of the solution of (5) with initial condition $x(t_0) = x_0$.

**Property 11.** (solutions of differential equations depend continuously on initial conditions) *Function $\phi_t$ is continuous.*

Similar results may be obtained under less demanding assumptions, allowing for generalizations of our results under similarly less demanding assumptions on the regularity of treatment level and of the sensitive population size.

2.5.2. *Proof of Proposition 1.* Assume that conditions i) and ii) hold, and then distinguish three cases.

**Case 1: if iiia) holds.** Let $f(t,R) = Rg_r(S_1(t), R)$, so that $\dot{R}_1(t) = f(t, R_1(t))$. Since $S_1 \geq S_2$ and $g_r$ is decreasing in $S$, it follows that:

$$\dot{R}_2(t) = R_2(t)g_r(S_2(t), R_2(t)) \geq R_2(t)g_r(S_1(t), R_2(t)) = f(t, R_2(t))$$

Since $R_1(t_0) \leq R_2(t_0)$, the comparison principle implies that $R_1 \leq R_2$ on $[t_0, t_1]$

**Case 2: if iiib) holds.** The proof that $R_1 \leq R_2$ on $[t_0, t_1]$ is as in the proof of a) but with $f(t,R) = Rg_r(N_1(t) - R, R)$. Since by assumption, $N_1 \geq N_2$, it follows that $S_1 \leq S_2$ on $[t_0, t_1]$.

**Case 3: if iiic) holds.** The idea of the proof is as follows: in forward time, as long as $S_1 \geq S_2$, the comparison principle implies $R_1 \leq R_2$. Similarly, as long as $R_1 \leq R_2$, since we also have $C_1 \leq C_2$, the comparison principle implies $S_1 \geq S_2$. Thus, as long as we have one of the properties that we want, we have the other. However, there is a chicken and egg problem. To solve it, we slightly perturb initial conditions to make sure that both properties hold strictly initially, and then use the fact that solutions of differential equations depend continuously on initial conditions.

Let $\varepsilon > 0$. Let $(S_1^\varepsilon, R_1^\varepsilon)$ be solution of (3) for the same treatment $C(t) = C_1(t)$ as $(S_1, R_1)$, but with initial conditions $S_1^\varepsilon(t_0) = S_1(t_0) + \varepsilon > S_2(t_0)$, and $R_1^\varepsilon(t_0) = R_1(t_0) - \varepsilon < R_2(t_0)$. Let $\tau \in [t_0, t_1]$. A variant of case 1 leads to:

**Lemma 12.** *If $S_1^\varepsilon(t) \geq S_2(t)$ on $[t_0, \tau]$, then $R_1^\varepsilon(t) < R_2(t)$ on $[t_0, \tau]$.*

A similar argument, using that $g_s$ is non-increasing in $R$ and in $C$, implies the following lemma:

**Lemma 13.** *If $R_1^\varepsilon(t) \leq R_2(t)$ on $[t_0, \tau]$, then $S_1^\varepsilon(t) > S_2(t)$ on $[t_0, \tau]$.*

Putting both lemmas together, we obtain:

**Lemma 14.** *For all $t$ in $[t_0, t_1]$, $R_1^\varepsilon(t) < R_2(t)$ and $S_1^\varepsilon(t) > S_2(t)$.*

*Proof.* Otherwise there exists a first time $\tau$ in $[t_0, t_1]$ such that $R_1^\varepsilon(\tau) \geq R_2(\tau)$ or $S_1^\varepsilon(\tau) \leq S_2(\tau)$. But on $[0, \tau]$, $R_1^\varepsilon(t) \leq R_2(t)$ and $S_1^\varepsilon(t) \geq S_2(t)$. Thus, by Lemmas 12 and 13, $R_1^\varepsilon(\tau) < R_2(\tau)$ and $S_1^\varepsilon(\tau) > S_2(\tau)$. This contradicts the definition of $\tau$. □

Since solutions of differential equations depend continuously on initial conditions, it follows from Lemma 14 that, for any $t$ in $[t_0, t_1]$

$$R_1(t) = \lim_{\varepsilon \to 0} R_1^\varepsilon(t) \leq R_2(t)$$

and similarly $S_1(t) \geq S_2(t)$.

#### 3. Explicit formulas

We compute below, for various treatments and models, the time it takes for tumor size to become strictly larger than an arbitrary threshold $N^* \geq N_0$. This time is denoted by $t_{N^*}(treatment)$. Times to progression, to treatment failure, and survival times are obtained by taking $N^*$ equal to $N_0$, $N_{tol}$, and $N_{crit}$, respectively. Section 3.1 studies general density-dependent models, and Section 3.2 some frequency-dependent ones. Formulas for Gompertzian growth are given in Section 3.3. Treatments considered were defined in Section 2.1.

3.1. **The purely density-dependent case.** This section studies the particular case of Model (3) where $g_s(S,R,C) = g(N,C)$ and $g_r(S,R) = g(N,0)$, with $g$ decreasing both in $N$ and in $C$. Thus:

$$\text{(6)} \qquad \begin{cases} \dot{S}(t) &= S(t)g(N(t),C(t)) \quad ; \quad S(0) = S_0 \geq 0 \\ \dot{R}(t) &= R(t)g(N(t),0) \qquad ; \quad R(0) = R_0 > 0 \end{cases}$$

Henceforth, we let $g(N) := g(N,0)$. Note that there is no cost of resistance.

3.1.1. *No treatment, ideal MTD, delayed ideal MTD.* For an untreated of fully resistant tumor, $\dot{N} = Ng(N)$. This equation may be solved by separation of variables. The time it takes for tumor size to grow from $N_1$ to $N_2 > N_1$ is:

$$t_{N_1 \to N_2} = \int_{N_1}^{N_2} \frac{dN}{Ng(N)}$$

Supplementary Table 2 gives an explicit expression of $t_{N_1 \to N_2}$ for various tumor growth-models.

With the above notation, the time it takes for an untreated tumor to become larger than $N^*$ is:

$$t_{N^*}(noTreat) = t_{N_0 \to N^*} = \int_{N_0}^{N^*} \frac{dN}{Ng(N)}$$

Under ideal MTD, the tumor is first reduced to size $R_0$ and then grows as an untreated tumor. Thus:

$$t_{N^*}(idMTD) = t_{R_0 \to N^*} = t_{N^*}(noTreat) + t_{R_0 \to N_0}$$

Under delayed ideal MTD, with treatment starting at some size $N_{ref} \geq N_0$, the tumor first grows to size $N_{ref}$, then is reduced to the current resistant population size $R_1$, and then grows back as a fully resistant tumor. Due to the absence of resistance cost, the frequency of resistant cells does not change during no treatment phases, so that $R_1 = R_0 \frac{N_{ref}}{N_0} \geq R_0$. For $N^* \geq N_{ref} \geq N_0$, this leads to:

$$t_{N^*}(del\text{-}idMTD) = t_{N_0 \to N_{ref}} + t_{R_1 \to N^*} = t_{N^*}(noTreat) + t_{R_0 N_{ref}/N_0 \to N_{ref}}$$

3.1.2. *Ideal containment at* $N_{ref} \geq N_0$. The tumor grows as an untreated or fully resistant tumor before and after the stabilization phase. For $N^* < N_{ref}$, $t_{N^*}(idCont) = t_{N^*}(noTreat)$. For $N^* \geq N_{ref}$, the absolute benefit of ideal containment with respect to no treatment is the duration of the stabilization phase. This is the time it takes for the resistant population to grow from $R_1 = R_0 \frac{N_{ref}}{N_0}$ to $R_2 = N_{ref}$ at a constant per-cell growth-rate $g(N_{ref})$. This leads to:

$$t_{N^*}(idCont) = t_{N^*}(noTreat) + \frac{\ln(N_0/R_0)}{g(N_{ref})}$$

3.1.3. *Constant dose, MTD, and delayed constant dose with a Norton-Simon kill rate.* Assume a Norton-Simon kill-rate: $g(N,C) = g(N)(1-\lambda C)$. Then under a constant dose treatment, $\frac{dS}{S} = (1-\lambda C)\frac{dR}{R}$ so that $SR^{\lambda C - 1}$ is constant. Thus, $S = S_0 \left(\frac{R_0}{R}\right)^{\lambda C - 1}$, and when $N = N^*$ for the last time, the resistant

Supplementary table 2. **Time in which an untreated tumor grows from $N_1$ to $N_2$.**

| Model name | Per-cell growth-rate $g(N)$ | $t_{N_1 \to N_2}$ |
|---|---|---|
| Exponential | $\rho$ | $\frac{1}{\rho} \ln\left(\frac{N_2}{N_1}\right)$ |
| Gompertz | $\rho \ln(K/N)$ | $\frac{1}{\rho} \ln\left(\frac{\ln(K/N_1)}{\ln(K/N_2)}\right)$ |
| Logistic | $\rho(1 - N/K)$ | $\frac{1}{\rho}\left[\ln\left(\frac{N_2}{N_1}\right) + \ln\left(\frac{K-N_1}{K-N_2}\right)\right]$ |
| Power-Law | $\rho N^{-\gamma}$; $0 < \gamma < 1$ | $\frac{1}{\rho\gamma}(N_2^\gamma - N_1^\gamma)$ |
| von Bertalanffy | $\rho(N^{-\gamma} - K^{-\gamma})$; $0 < \gamma < 1$ | $\frac{1}{\rho\gamma K^{-\gamma}} \ln\left(\frac{K^\gamma - N_1^\gamma}{K^\gamma - N_2^\gamma}\right)$ |

population size is the largest solution $R^*$ of:[8]

$$(7) \qquad N(R^*) = N^*,\ R^* \geq R_0,\ \text{with } N(R) = R + S_0\left(\frac{R_0}{R}\right)^{\lambda C - 1}$$

This leads to:

$$t_{N^*}(\text{Constant dose } C) = \int_{R_0}^{R^*} \frac{dR}{Rg(N(R))}$$

with $R^*$ and $N(R)$ defined by (7). A formula for MTD is obtained by taking $C = C_{max}$.

More generally, if at some point $N = N_1$, $R = R_1$, and tumor is then treated as a constant dose $C$, then the time it takes for the tumor to exceed size $N_2 \geq N_1$ is:

$$(8) \qquad t(N_1 \to N_2|R_1, C) = \int_{R_1}^{R_2} \frac{dR}{Rg(\tilde{N}(R))} \text{ with } \tilde{N}(R) = R + S_1\left(\frac{R_1}{R}\right)^{\lambda C - 1}$$

where $R_2$ is the largest solution of $R_2 + S_1\left(\frac{R_1}{R_2}\right)^{\lambda C-1} = N_2$. However, except in very special cases, $R_2$ and the integral can only be computed numerically.

If treatment is delayed until $N = N_{ref}$ and a constant dose $C$ is then applied, the time at which tumor size increases beyond $N^*$, assuming $N^* \geq N_{ref}$, is equal to

$$t_{N^*}(\text{delayed constant dose } C) = t_{N_0 \to N_{ref}} + t(N_{ref} \to N^*|R_1, C),\ \text{with } R_1 = R_0 N_0/N_{ref}$$

where the second term is defined by (8). Taking $C = C_{max}$ gives a formula for delayed MTD, though again with an integral to be computed numerically.

3.1.4. *Containment at $N_{ref}$*. If $N^* < N_{ref}$, then $t_{N^*}(Cont) = t_{N^*}(noTreat)$. If $N^* = N_{ref}$, then as for ideal containment:

$$t_{N_{ref}}(Cont) = t_{N_{ref}}(noTreat) + \frac{\ln(R_2/R_1)}{g(N_{ref})}$$

where $R_1 = R_0\frac{N_{ref}}{N_0}$ and $R_2$ are the resistant population sizes at the beginning and at the end of the stabilization phase. However, $R_2$ is no longer equal to $N_{ref}$, and needs to be computed. To do so, let $\tilde{R}_2$ be the solution of:

$$(N_{ref} - R)g(N_{ref}, C_{max}) + Rg(N_{ref}) = 0$$

that is, if $N = N_{ref}$, $R = \tilde{R}_2$ and $C = C_{max}$, then $dN/dt = 0$. There are two cases: if $R_1 \geq \tilde{R}_2$, or equivalently $R_0 \geq \frac{N_0}{N_{ref}}\tilde{R}_2$, then when the tumor reaches the stabilization size $N_{ref}$, treating at $C_{max}$ does not decrease tumor size: there is then no stabilization phase, $R_2 = R_1$ and $t_{N^*}(Cont) = t_{N^*}(noTreat)$. Otherwise, there is a stabilization phase that lasts until $R = \tilde{R}_2$. Thus, $R_2 = \max(R_1, \tilde{R}_2)$. In the case $R_1 < \tilde{R}_2$ (existence of a stabilization phase), we get:

$$t_{N_{ref}}(Cont) = t_{N_{ref}}(noTreat) + \frac{\ln(N_0/R_0)}{g(N_{ref})} - \frac{\ln\left(1 + g(N_{ref})/|g(N_{ref}, C_{max})|\right)}{g(N_{ref})}$$

For a Norton-Simon kill rate: $g(N, C) = g(N)(1 - \lambda C)$, this boils down to:

$$t_{N_{ref}}(Cont) = t_{N_{ref}}(noTreat) + \frac{\ln(N_0/R_0)}{g(N_{ref})} - \frac{\ln\left(\lambda C_{max}/[\lambda C_{max} - 1]\right)}{g(N_{ref})}$$

For $N^* \geq N_{ref}$, still assuming a Norton-Simon kill rate and the existence of a stabilization phase,

$$t_{N^*}(Cont) = t_{N_{ref}}(noTreat) + \frac{\ln(N_0/R_0)}{g(N_{ref})} - \frac{\ln\left(\lambda C_{max}/[\lambda C_{max} - 1]\right)}{g(N_{ref})} + t(N_{ref} \to N^*|\tilde{R}_2, C_{max})$$

where the last term is defined by (8). The difference with ideal containment is:

$$t_{N^*}(Cont) - t_{N^*}(idCont) = -\frac{\ln\left(\lambda C_{max}/[\lambda C_{max} - 1]\right)}{g(N_{ref})} + \left[t(N_{ref} \to N^*|\tilde{R}_2, C_{max}) - t_{N_{ref} \to N^*}\right]$$

The first term is the difference in the durations of the stabilization phases. It is smaller if $C_{max}$ is large. The second term (the bracket) is the difference between the time it takes for the tumor to progress from $N_{ref}$ to $N^*$ under containment and under ideal containment. It is positive and increasing in $N^*$. This expresses

[8] The solution to (7) is easily seen to be unique unless both $N^* = N_0$ and $S_0(1 - \lambda C) + R_0 < 0$ (i.e. tumor size initially decreases), in which case $R^*$ is the unique solution strictly greater than $R_0$.

the fact that, under containment, after the stabilization phase, the tumor is still partially sensitive, hence progresses more slowly than under ideal containment.

3.1.5. *Ideal intermittent containment between $N_{min}$ and $N_{max}$.* To fix ideas, assume $N_0 \leq N_{max} \leq N^*$. Under ideal intermittent containment, the tumor grows as an untreated or fully resistant tumor, except that each time it reaches size $N_{max}$ and is still partially sensitive the sensitive population is decreased by $N_{max} - N_{min}$, or by $N_{max} - R$ if $R > N_{min}$. Letting $t_{stab}(idInt)$ be the duration of the (dynamic) stabilization phase, that is, the time between the first and the last time such that $N = N_{max}$,

$$t_{N^*}(idInt) = t_{N^*}(noTreat) + t_{stab}(idInt)$$

We now compute $t_{stab}$. Let $t_k$ denote the $k$th time that $N = N_{max}$. Let $t_{q+1}$ denote the first time at which $N = N_{max}$ and $N$ cannot be reduced to $N_{min}$ (that is, $R > N_{min}$). Let $R_{q+1} = R(t_{q+1})$. During the stabilization phase, the tumor size changes $q$ times from $N_{min}$ to $N_{max}$, and once from $R_{q+1}$ to $N_{max}$. Therefore:

$$t_{stab} = q\, t_{N_{min} \to N_{max}} + t_{R_{q+1} \to N_{max}}$$

It remains to compute $q$ and $R_{q+1}$. Due to the absence of resistance cost, the proportion of resistant cells does not change during a no-treatment phase. This implies that:

$$R_{q+1} = R_0 \times \frac{N_{max}}{N_0} \times \left(\frac{N_{max}}{N_{min}}\right)^q \tag{9}$$

so it only remains to compute $q$. The fact that, by definition of $q$, $N_{min} < R_{q+1} \leq N_{max}$ implies that $q$ is the integer part of (i.e., the greatest integer no greater than)

$$\frac{\ln(N_0/R_0)}{\ln(N_{max}/N_{min})} \tag{10}$$

It may be checked that if $N_{min} < R_0$, so that $q = 0$, we obtain the formula for delayed ideal MTD (let the tumor grow until $N_{max}$, then eliminate all sensitive cells). Similarly, in the limit $N_{min} \to N_{max}$, we recover the formula for ideal (continuous) containment.

3.1.6. *Summary.* Supplementary Table 3 summarizes absolute benefits of various treatments compared to not treating:

$$t_{N^*}(treatment) - t_{N^*}(noTreat),$$

in the case $N^* \geq N_{ref} \geq N_0$. The formula for containment is given in the case of a Norton-Simon kill rate, and assuming that some stabilization is possible (for other cases, see Section 3.1.4). The inequality in this formula indicates that the benefit is greater that this quantity (with equality for $N^* = N_{ref}$). In simulations, for large values of $N^*$, we find the benefit of containment to be similar to the benefit of ideal containment, and even slightly greater in some cases when the endpoint tumor size is larger than the containment size.

3.2. **Frequency-dependent models and models with resistance costs.** Models where the resistant population follows a frequency-dependent dynamic $\dot{R} = Rf(R/N)$, or more generally a frequency and density-dependent dynamic,

$$\dot{R} = Rf(R/N)g(N), \text{ with } f \text{ increasing and } f(1) = 1 \tag{11}$$

have been considered in the literature, e.g., Silva *et al.* (2012) [13], Bacevic *et al.* (2017) [14]. Another possibility is to keep an essentially density dependent model, but to introduce a resistance cost, e.g., $\dot{S} = \rho_s S g(N)$, $\dot{R} = \rho_r R g(N)$, with $\rho_r \leq \rho_s$. Under such models, the frequencies of resistant and sensitive cells continuously change even in an untreated tumor. This makes it difficult to obtain useful explicit formulas for treatments that begin by letting the tumor grow, such as containment at tumor sizes higher than the initial size. But for ideal MTD and ideal containment at the initial size $N_0$, explicit formulas are

Supplementary table 3. **Absolute benefits compared to not treating in Model** (6)

| Treatment | idMTD | del-idMTD | idCont | Cont |
|---|---|---|---|---|
| Absolute benefit | $t_{R_0 \to N_0}$ | $t_{R_0 \frac{N_{ref}}{N_0} \to N_{ref}}$ | $\frac{\ln(N_0/R_0)}{g(N_{ref})}$ | $\geq \frac{\ln(N_0/R_0) - \ln(1 + 1/[\lambda C_{max} - 1])}{g(N_{ref})}$ |

readily obtained for general models where $\dot{R} = Rg_r(R,S)$. Indeed, under ideal MTD, the tumor becomes immediately fully resistant, hence the frequency-dependence disappears. Thus, for $N^* \geq N_0$, we still have

$$t_{N^*}(idMTD) = t_{R_0 \to N^*}, \text{ with } t_{N_1 \to N_2} = \int_{N_1}^{N_2} \frac{dN}{Ng(N)} \text{ for Model (11),}$$

or more generally $t_{N_1 \to N_2} = \int_{N_1}^{N_2} \frac{dN}{Ng_r(N,0)}$.

For ideal containment at $N_0$, the stabilization phase lasts:

$$t_{stab}(idContN_0) = \frac{1}{g(N_0)} \int_{R_0}^{N_0} \frac{dR}{Rf(R/N_0)} \text{ for Model (11),}$$

and more generally, $t_{stab}(idContN_0) = \int_{R_0}^{N_0} \frac{dR}{Rg_r(R,N_0-R)}$. After the stabilization phase, the tumor is fully resistant. Thus,

$$t_{N^*}(idContN_0) = t_{stab}(idContN_0) + t_{N_0 \to N^*}.$$

For containment at the initial size $N_0$, the stabilization phase lasts:

$$t_{stab}(ContN_0) = \frac{1}{g(N_0)} \int_{R_0}^{R_{end}} \frac{dR}{Rf(R/N_0)} \text{ for Model (11),}$$

and more generally, $t_{stab}(ContN_0) = \int_{R_0}^{R_{end}} \frac{dR}{Rg_r(R,N_0-R)}$, where $R_{end}$ is the resistant population size when the stabilization treatment level reaches $C_{max}$. The value of $R_{end}$ is easy to compute once the sensitive population dynamics are specified. For $N^* \geq N_0$,

$$t_{N^*}(ContN_0) \geq t_{stab}(ContN_0) + t_{N_0 \to N^*}.$$

The expression for $t_{N_1 \to N_2}$ no longer corresponds to the growth of an untreated tumor, but to the growth of a fully resistant tumor. Thus, the formulas do not permit easy comparison with no treatment, but they allow comparison of ideal MTD and ideal containment (or containment). For the same function $g(N)$ as in Section 3.1, frequency-dependence as modeled in (11) provides an additional rationale to containment, increasing its benefit with respect to ideal MTD.

3.3. **Application to Gompertzian growth.** We now consider the case of a Gompertz model: $g(N) = \rho \ln(K/N)$, for which:

$$t_{N_1 \to N_2} = \frac{1}{\rho} \ln \left[ \frac{\ln(K/N_1)}{\ln(K/N_2)} \right] = \frac{1}{\rho} \ln \left[ \frac{\log(K/N_1)}{\log(K/N_2)} \right]$$

where ln denotes the natural logarithm ($\ln(2.718...) = 1$), while log denotes the base 10 logarithm ($\log(10) = 1$).For readability, time units are chosen so that $\rho = 1$ (otherwise, all times should be divided by $\rho$). Assume $N_0 \leq N_{tol} \leq N_{crit} \leq K$, and let $a$, $b$, $c$, $d$ be nonnegative real numbers such that

$$K = 10^a R_0 = 10^b N_0 = 10^c N_{tol} = 10^d N_{crit}.$$

Note that $a \geq b \geq c \geq d$. For ideal intermittent containment, let $N_{max} = 10^\alpha N_{min}$. *With time units such that* $\rho = 1$, the time to progression, the time to treatment failure, and the survival time in the absence of treatment are respectively 0, $\ln(b/c)$ and $\ln(b/d)$. Supplementary Table 4 gives the *absolute benefit* compared to no treatment in terms of time to progression, time to treatment failure, and survival time. The formulas for ideal intermittent containment are approximations (exact when the last cycle is complete), see Section 3.1.5. Formulas for other models are easily obtained by using the values of $g(N)$ and $t_{N_1 \to N_2}$ in Supplementary Table 2.

Supplementary Tables 5 and 6 give numerical values of time to progression, time to treatment failure, and survival time with parameters from Monro and Gaffney (2009) [11] (main text Table 2).[9] Note that contrary to Supplementary Table 4, these are not the benefits with respect to no treatment, but the actual times to progression, time to treatment failure and survival time. This is why we add a row for no treatment in Supplementary Table 6. Note also that, although it leads in our model to a large survival time, we do not advocate containment at the critical size $N_{crit}$ as such a strategy would be extremely risky and harmful to quality of life.

[9] In [11], simulations start with $S = 1$, $R = 0$, so the value of $R$ at treatment initiation, i.e., when $N = 10^{10}$, is not explicitly given. However, it may be derived from a well-known formula, see, e.g., Goldie and Coldman, 1979 [20].

Supplementary table 4. **Absolute benefit over no treatment for a Gompertz model: formulas.** Parameters satisfy $K = 10^a R_0 = 10^b N_0 = 10^c N_{tol} = 10^d N_{crit}$, and $N_{max} = 10^\alpha N_{min}$. Time units exceptionally chosen so that $\rho = 1$ (otherwise all times should be divided by $\rho$). Time to progression, to treatment failure, and survival time in the absence of treatment are respectively 0, $\ln(b/c)$ and $\ln(b/d)$, which should be added to the values of the table to obtain the correspond times for the treatment of interest.

| Treatment | Progression benefit | Treatment failure benefit | Survival benefit |
|---|---|---|---|
| ideal MTD | $\ln\left(1+\frac{a-b}{b}\right)$ | $\ln\left(1+\frac{a-b}{b}\right)$ | $\ln\left(1+\frac{a-b}{b}\right)$ |
| del-idMTD $N_{ref} = N_{tol}$ | 0 | $\ln\left(1+\frac{a-b}{c}\right)$ | $\ln\left(1+\frac{a-b}{c}\right)$ |
| del-idMTD $N_{ref} = N_{crit}$ | 0 | 0 | $\ln\left(1+\frac{a-b}{d}\right)$ |
| idCont$N_0$ | $\frac{a-b}{b}$ | $\frac{a-b}{b}$ | $\frac{a-b}{b}$ |
| idCont$N_{tol}$ | 0 | $\frac{a-b}{c}$ | $\frac{a-b}{c}$ |
| idCont$N_{crit}$ | 0 | 0 | $\frac{a-b}{d}$ |
| idInt$N_0$ | $\frac{a-b}{b} \times \frac{\ln(1+\alpha/b)}{\alpha/b}$ | $\frac{a-b}{b} \times \frac{\ln(1+\alpha/b)}{\alpha/b}$ | $\frac{a-b}{b} \times \frac{\ln(1+\alpha/b)}{\alpha/b}$ |
| idInt$N_{tol}$ | 0 | $\frac{a-b}{c} \times \frac{\ln(1+\alpha/c)}{\alpha/c}$ | $\frac{a-b}{c} \times \frac{\ln(1+\alpha/c)}{\alpha/c}$ |
| idInt$N_{crit}$ | 0 | 0 | $\frac{a-b}{d} \times \frac{\ln(1+\alpha/d)}{\alpha/d}$ |
| Cont$N_0$ | $\frac{a-b-\log\left(\frac{\lambda C_{max}}{\lambda C_{max}-1}\right)}{b}$ | no formula | no formula |
| Cont$N_{tol}$ | 0 | $\frac{a-b-\log\left(\frac{\lambda C_{max}}{\lambda C_{max}-1}\right)}{c}$ | no formula |
| Cont$N_{crit}$ | 0 | 0 | $\frac{a-b-\log\left(\frac{\lambda C_{max}}{\lambda C_{max}-1}\right)}{d}$ |

Supplementary table 5. **Times to progression, to treatment failure, and survival time for a Gompertz model: idealized treatments and containment** The model is Model 2 from the main text, with parameters from Table 1. Time is measured in days. Numbers with an asterisk are estimated from simulations, others are calculated from formulas.

| Treatment | $t_{prog}$ | $t_{fail}$ | $t_{surv}$ |
|---|---|---|---|
| ideal MTD | 186 | 263 | 412 |
| delayed ideal MTD (at $N_{tol}$) | 0 | 319 | 468 |
| delayed ideal MTD (at $N_{crit}$) | 0 | 77 | 591 |
| ideal containment at $N_0$ | 340 | 417 | 566 |
| ideal containment at $N_{tol}$ | 0 | 615 | 764 |
| ideal containment at $N_{crit}$ | 0 | 77 | 1526 |
| idInt between $N_0$ and $N_0/2$ | 320 | 397 | 546 |
| idInt between $N_{tol}$ and $N_{tol}/2$ | 0 | 566 | 715 |
| idInt between $N_{crit}$ and $N_{crit}/2$ | 0 | 77 | 1280 |
| Containment at $N_0$ ($C_{max} = 2$) | 318 | 418* | 568* |
| Containment at $N_{tol}$ ($C_{max} = 2$) | 0 | 580 | 767* |
| Containment at $N_{crit}$ ($C_{max} = 2$) | 0 | 77 | 1441 |

Supplementary table 6. **Times to progression, to treatment failure, and survival time for a Gompertz model: constant doses and delayed constant doses** Same model and parameters as in Supplementary Table 5. Numbers in parentheses (given only when different) are for the original Monro and Gaffney model [11], with mutations and back mutations at rate $10^{-6}$ (this does not change optimal doses, given our precision level). Bold squares correspond to optimal results given the starting time, e.g., the dose $C = 0.74$ maximizes survival time among all constant dose treatments starting immediately. $C = 2$ corresponds to MTD in most of our simulations. For visual representation of survival time for constant doses starting immediately, see Figure 1 in [11].

| C | Starting size | $t_{prog}$ | $t_{fail}$ | $t_{surv}$ |
|---|---|---|---|---|
| 2 (MTD) | $N_0$ | 236 | 314 | 463 |
| 1.09 | $N_0$ | **303 (302)** | 397 (396) | 549 (548) |
| 1 | $N_0$ | 0 | 421 (420) | 578 (579) |
| 0.89 | $N_0$ | 0 | **443 (442)** | 634 (632) |
| 0.74 | $N_0$ | 0 | 296 | **730 (727)** |
| 0 | irrelevant | 0 | 77 | 226 |
| 2 (MTD) | $N_{tol}$ | 0 | 400 (398) | 549 (547) |
| 1.07 | $N_{tol}$ | 0 | **543 (537)** | 731 (726) |
| 1 | $N_{tol}$ | 0 | 77 | 780 (774) |
| 0.86 | $N_{tol}$ | 0 | 77 | **885 (877)** |
| 2 (MTD) | $N_{crit}$ | 0 | 77 | 762 (758) |
| 1.04 | $N_{crit}$ | 0 | 77 | **1276 (1255)** |

#### 4. Comparison between treatments

Building on findings of Sections 2 and 3, this section compares treatments for a purely density-dependent model with no resistance cost, as defined in Section 3.1, Eq. (6).

##### 4.1. Idealized treatments.

4.1.1. *Impact of tumor growth model.* To fix ideas, assume $N^* \geq N_{ref} = N_{max} \geq N_0$. Recall that $t_{N^*}(treatment)$ denote the time at which tumor size exceeds size $N^*$ under the treatment considered. It follows from Proposition 7 that already in the general model of Section 2,

$$t_{N^*}(idMTD) \leq t_{N^*}(del\text{-}idMTD) \leq t_{N^*}(idInt) \leq t_{N^*}(idCont) \tag{12}$$

Considering a purely density-dependent model leads to further insights. In this case, the benefits of ideal MTD, delayed ideal MTD, ideal intermittent containment, and ideal containment, compared to not treating, are each the time taken for the resistant population to grow by a factor $N_0/R_0$, but in different circumstances: from $R_0$ to $N_0$ in the absence of competition for ideal MTD; from $R_1 = \frac{R_0}{N_0}N_{ref}$ to $N_{ref}$ with no, some, or strong competition for delayed ideal MTD, ideal intermittent containment, and ideal containment, respectively. Since doubling times are assumed longer at higher tumor sizes, we obtain another proof of (12). Moreover, the duration of the stabilization phase is inversely proportional to $g(N_{ref})$, hence the larger the stabilization size, the longer the stabilization phase, and the greater the benefits of ideal containment. (Figs. 1f, 1h; Supplementary Tables 3, 4, 5).

These qualitative findings are very general. However, the magnitude of clinical benefits vary considerably depending on the precise model [6, 7]. As easily seen, a general bound on the relative benefits of ideal containment over ideal MTD in terms of time to progression is given by:

$$\frac{t_{prog}(idContN_0)}{t_{prog}(idMTD)} \leq \frac{g(R_0)}{g(N_{ref})}$$

For logistic growth, $g(N) = 1 - N/K$, so that:

$$\frac{t_{prog}(idContN_0)}{t_{prog}(idMTD)} \leq \frac{1 - R_0/K}{1 - N_0/K} \leq \frac{1}{1 - N_0/K} \simeq 1 + \frac{N_0}{K}$$

for $N_0/K$ small. With the parameters from main text Table 2, $N_0/K = 1/200$, so that ideal containment improves on ideal MTD by less than 0.5%, a very tiny gain. To take into account the fact that the value of the carrying capacity $K$ was chosen for a Gompertz model [11], we may want to modify this value (see

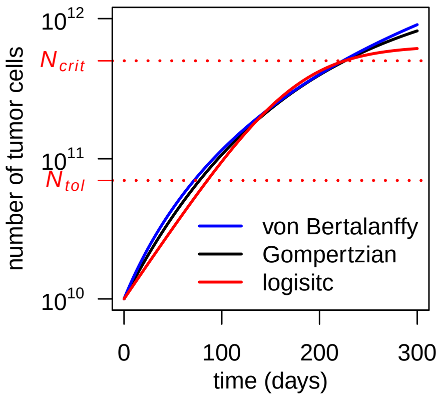


Supplementary figure 1. **Tumor growth-curves for three models with different forms of density dependence.** Untreated tumor growth curves for a Gompertzian growth model (black curve; Model 2 in the main text), a logistic growth model (red) and a von Bertalanffy growth model (blue) (these models are defined in Supplementary Table 2). Parameter values for the Gompertzian growth model are as in main text Table 1. Parameter values of the other two models are chosen so that the three curves are similar for tumor sizes between $N_0$ and $N_{crit}$ (the lethal size), as would be the case if the models were fitted to empirical data. In the logistic growth model, $K = 6.4 \times 10^{11}$ and $\rho = 2.4 \times 10^{-2}$. In the von Bertalanffy growth model, $K = 5 \times 10^{13}$, $\rho = 90$ and $\gamma = 1/3$ (the latter value is conventional in tumor growth modeling [21, 22]).

Supplementary Fig. 1). But as long as $K \geq N_{crit}$, and $N_{crit} = 50N_0$ as in main text Table 2, we still have $N_0/K \leq 0.02$ so that ideal containment improves on ideal MTD by less than about 2%. By contrast, with parameters from Table 2 but a Gompertz model, the time to progression under ideal containment at the initial size is 84% higher than under ideal MTD (Supplementary Table 5). Moreover, relative benefits of ideal containment for a von Bertalanffy model are even higher than for a Gompertz model (Fig. 2c). Thus, these three standard tumor growth models lead to very different quantitative predictions. We plan to study the influence of the model on predicted benefits of containment in a companion paper.

4.1.2. *Impact of parameters.* We discuss here the effect of varying parameters on absolute and relative benefits of ideal containment over ideal MTD, for the Gompertzian Model 2 of the main text. Changing the baseline growth-rate $\rho$ just changes the time-scale. Halving $\rho$ doubles absolute differences between treatments, hence absolute benefits of ideal containment, but does not change relative benefits. To investigate the effect of other parameters, let $x = \ln(K/R_0)/\ln(K/N_0)$. As follows from Section 3, the absolute benefit of ideal containment at $N_0$, and its relative benefit in terms of time to progression, are given by $x - \ln(1 + x)$ and $x/\ln(1 + x)$, respectively (Supplementary Table 4). These are two increasing functions of $x$, so any change of parameter that increases the value of $x$ increases both of these benefits. It is easy to see that this is the case of a decrease in $K$ or in $R_0$ (with $N_0$ fixed), or of an increase in $N_0$ (with $R_0$ fixed, and also with $R_0/N_0$ fixed). The absolute benefit of ideal containment at $N_{tol}$ with respect to ideal MTD is also easy to study. Results are summed up in Supplementary Table 7.

Supplementary Fig. 2 illustrates the impact of increasing the initial fraction of resistant cells on times to progression in Model 2 of the main text (see also Fig. 2 and Supplementary Fig. 3). For a given average per-cell growth rate, increasing $R_0$ reduces the time it takes for the resistant population to increase from $R_0$ to $N_0$. This tends to decrease absolute difference between treatments. Moreover, while this does not change the per-cell growth rate of resistant cells during the stabilization phase of containment or ideal containment, increasing $R_0$ increases the per-cell growth rate during the regrowth phase of MTD or ideal MTD, so that the differences in growth rates is smaller. This is because a higher initial resistant population leads to a higher average tumor size during this regrowth phase. This further reduces absolute benefits of ideal containment, and also reduces relative benefits.

Supplementary table 7. **Qualitative effect of parameters on absolute benefits of ideal containment over ideal MTD in main text Model 2.** The table should be read as follows: a "+" means an increase, an "=" no change, and a "-" a decrease.

| Parameter increased | $K$ | $N_{tol}$ | $N_0$ ($R_0$ fixed) | $N_0$ ($N_0/R_0$ fixed) | $R_0$ ($N_0$ fixed) |
|---|---|---|---|---|---|
| Effect on $t_{prog}(idContN0) - t_{prog}(idMTD)$ | - | = | + | + | - |
| Effect on $t_{fail}(idContN_{tol}) - t_{fail}(idMTD)$ | - | + | + | - | - |

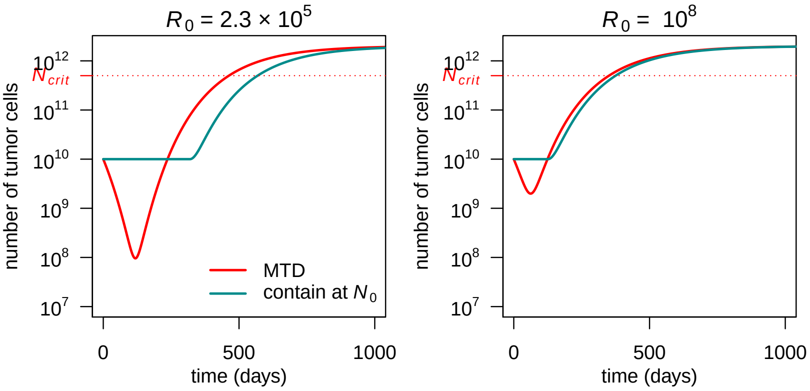


Supplementary figure 2. **Evolution of total tumor size under containment and MTD treatment in a Gompertzian growth model (Model 2 in the main text).** The initial resistant subpopulation size ($R_0$) is varied. The maximum dose is $C_{max} = 2$. Fixed parameter values are as in Table 1 of the main text.

The impact of varying parameters on *relative benefits* in terms of *time to treatment failure* or survival time is more complex. Indeed, varying a parameter may increase the absolute benefit of ideal containment over ideal MTD, yet reduce its relative benefit. This may happen for instance if it also increases the duration of growth phases that are common to ideal containment and ideal MTD. Indeed, a long common phase attenuates relative differences between treatments.

Consider for instance the impact of a lower initial tumor size $N_0$ for a fixed initial fraction of resistant cells $R_0/N_0$. Assuming $N_0 < N_{tol}$, this does not change the duration of the stabilization phase under ideal containment at $N_{tol}$, but decreases the duration of the phase of regrowth from $R_0$ to $N_0$ under ideal MTD. As a result, this increases the absolute benefit of ideal containment at $N_{tol}$ over ideal MTD in terms of time to treatment failure. But this also increases the duration of the phase of growth from $N_0$ to $N_{tol}$, which is common to both treatments. The net effect on the ratio of times to treatment failure $t_{fail}(idContN_{tol})/t_{fail}(idMTD)$ is unclear. This is apparent from Fig. 2b in the main text, where we see that for a fixed initial fraction of resistant cells, a higher initial tumor size sometimes increases and sometimes decreases the relative benefit of ideal containment at $N_{tol}$.

4.2. **Containment and ideal containment.** As we saw in Section 2, Proposition 3, the tumor progresses beyond the stabilization size faster under containment than under ideal containment. However, the resistant population under containment is always smaller, leading intuitively to a survival time that is longer or at least comparable to that under ideal containment. For a purely density-dependent model, we may be more precise. In the case of a Norton-Simon kill rate, for instance, it follows from Section 3 that the ratio of the duration of the stabilization phases of containment and ideal containment is given by:

$$\frac{t_{stab}(Cont)}{t_{stab}(idCont)} = 1 + \frac{\ln(1 - 1/\lambda C_{max})}{\ln(N_0/R_0)}$$

(this ratio is the same independently of the stabilization size). Supplementary Table 8 gives this ratio for various initial proportions of resistant cells and efficiency $\lambda C_{max}$ of the MTD treatment (for a visual representation, see Fig. 2g in main text).[10] When sensitive cells are not very sensitive and resistant cells are initially abundant, this ratio is substantially smaller than 1. However, if the drug is quite effective or resistant cells are initially rare, this ratio is quite high. In that case, containment is not far from maximizing the time at which tumor size grows above the stabilization level (see also Figs. 1a, 1d, 2d, 2h, 2g, Supplementary Tables 4 and 5, Supplementary Fig. 3).[11]

Moreover, the reason why containment progresses beyond $N_{ref}$ before ideal containment is that this happens before all sensitive cells have been eliminated. With a Norton-Simon kill rate, this may be made precise: the proportion of sensitive cells when the tumor can no longer be stabilized (whatever the stabilization level) is either the initial proportion (when the tumor cannot be stabilized at all) or $1/\lambda C_{max}$: 50% if $\lambda C_{max} = 2$; 33% if $\lambda C_{max} = 3$; and so on.

These remaining sensitive cells slow down the expansion of resistant cells. This is why, as shown in Section 2, the resistant population is always smaller for containment than for ideal containment. A sufficient condition for containment to be at least comparable to ideal containment in terms of survival time is thus that, by the time tumor size reaches $N_{crit}$, almost all sensitive cells have been eliminated. This is more likely if the treatment is quite efficient against sensitive cells ($\lambda C_{max}$ high) and if the tumor is stabilized at a relatively low size, leaving more time after the end of the stabilization phase to eliminate the remaining sensitive cells. However, we now argue that even if containment is made at a relatively high tumor size, and treatment effect is relatively modest, it is likely that by the time of death, the proportion of sensitive cells will be very low.

To see this, consider the post-stabilization phase during which containment treats at $C_{max}$ and assume a Norton-Simon kill rate: $\dot{S} = Sg(N)(1 - \lambda C)$. As we saw in Section 3.1.3, if the tumor is treated at a constant dose $C$, then the quantity $SR^{\lambda C-1}$ is constant. It follows that if $\lambda C = 2$ (resp. 3), then each time the resistant population is multiplied by 10, the sensitive population is divided by 10 (resp. 100). Since we also know that the proportion of sensitive cells at the end of the stabilization phase is $1/\lambda C_{max}$, this allows us to estimate the remaining sensitive population at the time of death. For instance, assuming that containment is made at $N_{tol} = 7 \times 10^{10}$, that $N_{crit} = 5 \times 10^{11}$ (about 7 times larger than $N_{tol}$) and that $\lambda C_{max} = 2$, the proportion of sensitive cells at the time of death is only about 0.5%. This is true for any purely density-dependent model.

We conclude that in terms of survival time, containment is bound to be at least as good as ideal containment, and possibly slightly better. This is what we observe in simulations (Figs. 1a, 1c, 1d, 2f, 2i, Supplementary Table 5, and Supplementary Fig. 3).

4.3. **MTD and ideal MTD.** To facilitate the comparison between MTD and ideal MTD, let us first estimate the remaining sensitive population at the time of progression under the MTD treatment. Assume a Norton-Simon kill rate. Then, as discussed in Section 4.2, if $\lambda C_{max} = 2$ (resp. 3; 1.5), each time the

[10]If $\lambda C_{max} = 2$, the speed at which a purely sensitive tumor size decreases under MTD is equal to the speed at which it would increase if untreated. This is roughly consistent with the fact that, in preliminary results of a clinical trial of intermittent containment (Zhang et al, 2017 [1]), treatment was applied, on average, 47% of the time (this average later decreased to 41%, as stated by Robert A. Gatenby in a seminar on July 22, 2020). For this reason, and due to the simple interpretation it offers, we chose $\lambda C_{max} = 2$ as our reference value in simulations.

[11]If we take the constraint $C \leq C_{max}$ into account then containment is closer to optimal than the numbers seem to indicate. This is because the optimal strategy then begins as containment and switches to MTD before progression or failure occurs. Comparing containment to ideal containment gives us a non-tight upper bound on how far we are from optimality, given the constraint on $C_{max}$.

Supplementary table 8. **Ratio of times to progression under containment vs ideal containment for various initial fractions of resistant cells and treatment efficiencies.**

| | | $\lambda C_{max}$ | | |
|---|---|---|---|---|
| | | 1.5 | 2 | 5 |
| $\frac{R_0}{N_0}$ | 10% | 0.52 | 0.70 | 0.90 |
| | 1% | 0.76 | 0.85 | 0.95 |
| | 0.1% | 0.84 | 0.90 | 0.97 |
| | 0.01% | 0.88 | 0.92 | 0.96 |
| | 0.005% | 0.90 | 0.94 | 0.98 |

resistant population is multiplied by 10, the sensitive population is divided by 10 (resp. 100; $\sqrt{10} \simeq 3.2$). So if there were 1% resistant cells initially, then at time to progression, there will be 1% sensitive cells (resp. about 0.01%; about 10.5%). This shows that unless the initial proportion of resistant cells is very large, or treatment is very inefficient against sensitive cells, the MTD treatment will eliminate the vast majority of sensitive cells before progression.

Thus, the difference in times to progression mostly comes from the difference in resistant populations. Under the MTD treatment, the sensitive population disappears more slowly than under its idealized counterpart. As a consequence, the resistant population still somewhat competes with sensitive cells, and develops more slowly (Fig. 1c). The MTD treatment is thus expected to lead to a longer time to progression than its idealized version. This is especially true if treatment is relatively inefficient. In that case, by the time sensitive cells have been crushed, resistant cells are already abundant. It follows that tumor size is never very low, so that resistant cells never develop very quickly.

To quantify this phenomenon in the case of a Norton-Simon kill rate, recall that under MTD, the quantity $SR^{\lambda C_{max}}$ is constant. It follows that the tumor reaches its smallest size when $S = R/(\lambda C_{max} - 1)$. Its size is then:

$$N = \frac{\lambda C}{\lambda C - 1} \times \left[(\lambda C - 1) S_0 R_0^{\lambda C - 1}\right]^{1/\lambda C}, \text{ with } C = C_{max}$$

Some values of this minimal size are given in Supplementary Table 9, assuming $S_0 = 10^{10}$ and $R_0 = 10^6$. It confirms that, for modest treatment effects ($C_{max} \leq 2$), the minimal tumor size under MTD is much higher than under ideal MTD (that is, than the initial resistant population).

Finally, the fact that few sensitive cells remain at the time of progression implies that, after progression, MTD and ideal MTD have similar dynamics. As a result, the difference in survival times should not be much higher than the difference in times to progression. This is what we observe in simulations (Table 2, Figs. 2d, 2f, Supplementary Figs. 4 and 3)

4.4. **MTD and containment.** Ideal containment at $N_0$ always leads to a higher time to progression than ideal MTD, and with a Gompertz model, typically substantially so, unless the tumor is initially very resistant. The difference in time to progression between the more realistic versions – containment at $N_0$ and MTD – is smaller. If treatment is not very efficient, MTD may even lead to a higher time to progression than containment. For instance, in main text Model 2, if parameter values (other than $R_0$) are as in main text Table 1, this occurs whenever the initial fraction of resistant cells is higher than about 1% if $\lambda C_{max} = 2$ (Fig. 2d) and than about 0.1% if $\lambda C_{max} = 1.5$.

There are two explanations. First, low treatment efficiency decreases time to progression under containment. Indeed, progression then occurs as the tumor is still quite sensitive (see Fig. 2g, and Section 4.2). Second, as discussed in Section 4.3, low treatment efficiency makes MTD less problematic. Supplementary Table 9 shows that, for $\lambda C_{max} = 1.5$, the minimal tumor size under MTD is only slightly below $10^9$. The average size before progression will be substantially higher than $10^9$, compared to $N_0 = 10^{10} + 10^6 \simeq 10^{10}$ for containment at the initial size. In log-scale, this is not a huge difference. Thus, resistant cells would develop only slightly slower under containment than under MTD. With a higher initial resistant population, this advantage of containment is even lower, and need not compensate the fact that under containment, progression requires a lower resistant population size than under MTD, due to the larger remaining sensitive population.

However, if containment is made at a higher level, the average tumor burden during the stabilization phase is larger, and the benefit of containment over MTD in terms of the time to progress beyond size $N_{ref}$ is expected to be greater (compare Figs. 2d and 2h). Moreover, even when MTD and containment at $N_0$ are comparable in terms of time to progression, the resistant population is always smaller under containment, leading to longer times to treatment failure and longer survival times under containment than under MTD. This is seen in simulations (Figs.2d, 2e, 2f).

4.5. **Impact of varying $C_{max}$.** Supplementary Fig. 3 further explores the impact of varying the resistant population size $R_0$ and the maximal tolerated dose $C_{max}$ on containment at the maximal tolerable size,

Supplementary table 9. **Minimal tumor size under MTD.** These numbers are valid for any density-dependent model (6), assuming $S_0 = 10^{10}$ and $R_0 = 10^6$

| $\lambda C_{max}$ | 1.1 | 1.25 | 1.5 | 2 | 3 | 5 | 10 |
|---|---|---|---|---|---|---|---|
| Minimal N | $5.9 \times 10^9$ | $2.6 \times 10^9$ | $8.7 \times 10^8$ | $2 \times 10^8$ | $4.1 \times 10^7$ | $1.0 \times 10^7$ | $3.5 \times 10^6$ |

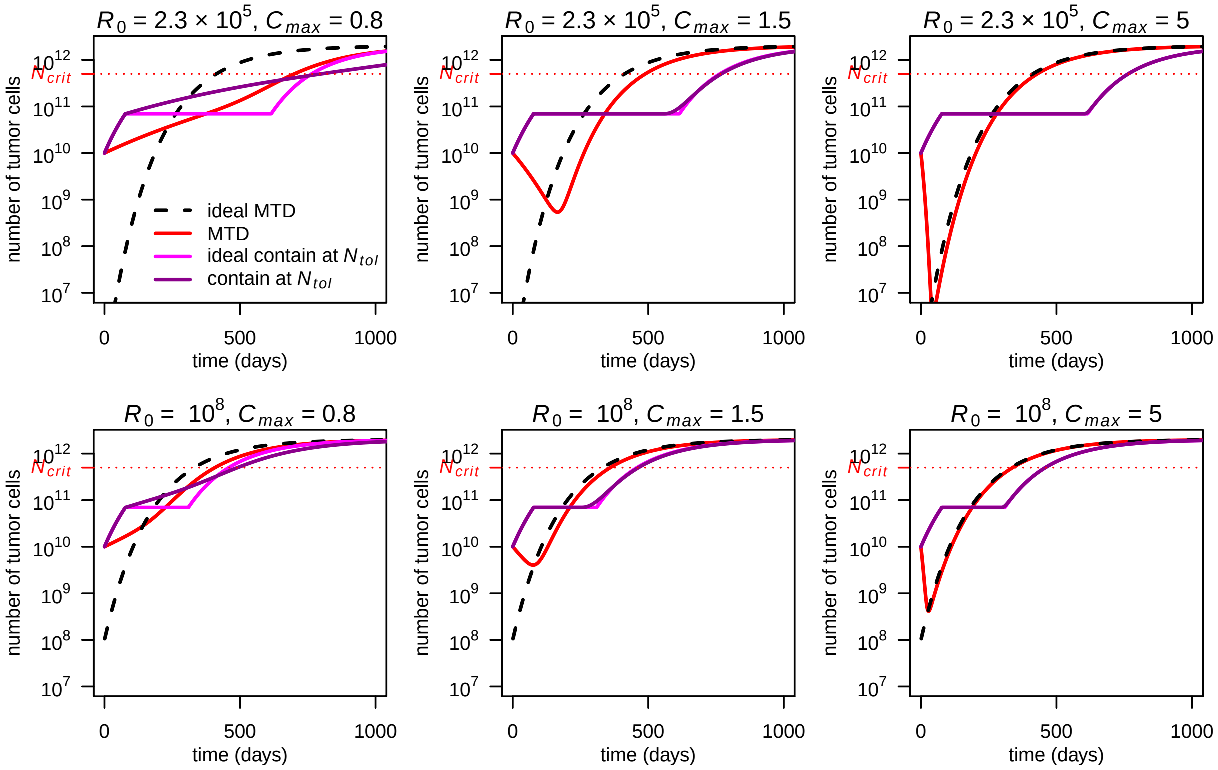


SUPPLEMENTARY FIGURE 3. **Evolution of total tumor size under ideal and non-ideal treatments in a Gompertzian growth model (Model 2 in the main text).** The initial resistant subpopulation size ($R_0$) and the maximum dose ($C_{max}$) are varied. Fixed parameter values are as in Table 1 of the main text.

MTD, and idealized versions. The doses considered have the following interpretation: If $C_{max} = 0.8$ then the sensitive cell population keeps growing under MTD, but 5 times slower than in the absence of treatment. If $C_{max} = 1.5$ (respectively 5) then the sensitive population decreases twice as slowly (respectively, four times faster) under MTD than it would increase in the absence of treatment.

In the first row, $R_0 = 2.3 \times 10^5$. If $C_{max} = 0.8$ (panel a) then there is no stabilization phase under containment. Containment at $N_{tol}$ then boils down to delayed MTD. The time to treatment failure is the same as under no treatment, and much lower than under MTD. Nevertheless, survival time is still higher than under MTD (though this would not be true for even lower values of $C_{max}$). Tumor composition at the time of death is very different between treatments: less than 5% of cells are resistant under containment at $N_{tol}$, versus 66% for MTD. Thus, under MTD, even though the sensitive population always grows, death occurs mostly due to resistant cells. Since tumor size is never reduced under MTD in this scenario, resistant cells develop much slower than under ideal MTD, leading to a much larger survival time.

If $C_{max} = 1.5$ (panel b), the sensitive population size decreases under MTD but relatively slowly. The minimal tumor size under MTD is much higher than under ideal MTD, leading to a substantial difference between the two treatments. Under containment, there is a stabilization phase that lasts until the tumor is 1/3 resistant. Given the low initial resistant population, there is little difference between growing from $R_0$ to $N_0/3$ and growing from $R_0$ to $N_0$, and so there is little difference between containment and ideal containment. Under all treatments, the tumor is almost fully resistant at the time of death (2% of cells are sensitive under containment, and 0.001% under MTD).

If $C_{max} = 5$ (panel c), the sensitive population decreases much faster than the resistant population increases, and there is very little difference between idealized and non-idealized treatments.

In the second row, $R_0 = 10^8$. Since the tumor is initially much more resistant, the tumor size never becomes very low, even under ideal MTD. As a result, there are smaller differences between treatments, but otherwise the impact of varying $C_{max}$ is similar. For $C_{max} = 0.8$, tumor composition at the time of death depends substantially on treatment: 89% of cells are resistant under MTD, but only 54% under containment at $N_{tol}$. For $C_{max} = 1.5$ and $C_{max} = 5$, the tumor is almost fully resistant by the time of death under all treatments.

4.6. **Constant dose and containment.** Survival time under constant dose and delayed constant dose treatments in main text Model 2 have been studied by Monro and Gaffney (2009) [11]. A key insight is that there is an optimal balance between limiting the expansion of sensitive cells, and keeping enough of them to slow down the growth of resistant cells. In other words, a tradeoff between dying from sensitive cells and dying from resistant cells. Constant doses that lead to high survival time are such that at the time of death, the populations of resistant and sensitive cells are of the same order of magnitude. Here we are also interested in time to progression and time to treatment failure, and in comparing with containment.

In Model 2, among constant dose treatments, survival time is maximized by the dose $C = 0.74$ (Supplementary Table 6 and Supplementary Fig. 4).[12] Being smaller than the dose used for containment (which is always greater than 1), the optimal constant dose leads to a smaller resistant population, while still significantly slowing down the growth of sensitive cells. This treatment strategy turns out to prolong survival more than either containment or ideal containment at the initial size $N_0$ (Supplementary Table 5). This does not violate our optimality results because ideal containment at $N_0$ maximizes time to progression, not survival time. The dose $C = 0.74$, however, is far from maximizing survival time among all possible treatments (Supplementary Table 5): treating even less initially represses resistant cells more efficiently, while treating more eventually prolongs survival by diminishing the sensitive population once this becomes necessary. Containment at a high threshold does both, leading to longer survival.

As discussed in the main text, containment may be mimicked by delaying treatment and then applying a dose $C = 1/\lambda$, or slightly higher. There are however two issues. First, as mentioned in the main text, the dose that allows stabilizing tumor size is bound to be patient-dependent. By adjusting the dose as a function of patient's response, containment allows the clinician to arrive at the right dose without foreknowledge. Second, treatment response, in practice, is likely to be much less predictable than our model suggests. In a pre-clinical trial, it was found that, possibly due to normalization of tumor vasculature, tumor control could be achieved by applying progressively lower doses [12]. In a clinical trial of intermittent containment

[12] According to our simulations, the dose that maximizes survival time among doses given immediately is $C = 0.74$ instead of the dose $C = 0.9$ reported by Monro and Gaffney (2009), which however is not consistent with the curves of their Fig. 1. This seems to be a simple typo: in Fig 1 of [11], point A visually seems to corresponds to $C \simeq 0.5$ and point $B$ (the optimum) to $C = 0.74$ as we find. There seem to be also typos in their description of Figs. 2 and 6. This does not change the study's key messages, which we agree with.

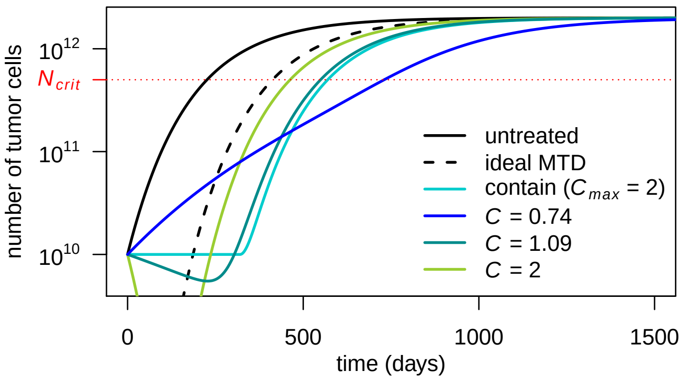


Supplementary figure 4. **Constant dose treatments in a Gompertzian growth model (Model 2 in the main text).** Tumor size for various constant dose treatments are compared to containment at the initial size (subject to $C_{max} = 2$), MTD ($C = C_{max}$) and ideal MTD. Dose 1.09 maximizes time to progression and 0.74 maximizes survival time among non-delayed constant doses (but is inferior to the optimal delayed constant dose). Parameter values are as in Table 1 (adaptive therapy) for patients suffering from metastatic castration resistant prostate cancer, tumor size cycled less regularly than models predicted [1]. This suggests that even for a given patient with known tumor characteristics, delayed constant doses might not mimic containment as well as our model suggests, and hence an adaptive protocol is necessary to even approximately stabilize tumor size.

#### 5. Impact of resistance costs on the best possible outcome and on clinical benefits of containment

5.1. **Justification of main text Figure 4b.** We discuss here the impact of competition and resistance costs on the best possible outcome for the following model (Model 3 in the main text):

$$\begin{aligned} \dot{S}(t) &= \rho_s \left[\ln\left(\frac{K_s}{S(t)+\alpha R(t)}\right)\right](1-\lambda C(t))S(t), \\ \dot{R}(t) &= \rho_r \ln\left(\frac{K_r}{R(t)+\beta S(t)}\right) R(t). \end{aligned}$$

If sensitive cells are eliminated by a high-dose treatment, then the resistant population size, hence tumor size, will grow to $K_r$, unless the patient dies before. Such a treatment would thus result in a tolerable long-run outcome (i.e. tumor size always below $N_{tol}$) only if $K_r \leq N_{tol}$. If $N_{tol} < K_r < N_{crit}$, the long-run tumor size would be non-lethal, but intolerable. If $K_r > N_{crit}$, a high dose treatment leads to death.

Whether containment strategies can do better depends on whether $\beta$ is greater than 1, that is, whether additional sensitive cells inhibit the growth of resistant cells more than additional resistant cells would. If $\beta \leq 1$, then $R + \beta S \leq R + S = N$. Thus, as long as $N < K_r$, $R + \beta S < K_r$, and the resistant population grows ($\dot{R} > 0$). In that case, containment strategies may gain time, but cannot change the long-run outcome.

If $\beta > 1$, then provided that the initial resistant population is low enough, the tumor could be stabilized around $S = K_r/\beta$, $R = 0$, hence a total tumor size of $N = K_r/\beta$. The tumor cannot be stabilized at a lower size. Indeed, if $N < K_r/\beta$, then since $\beta \geq 1$, $R + \beta S \leq \beta N < K_r$, hence the resistant population grows. The best long-run outcome is thus a tolerable tumor size if $K_r/\beta < N_{tol}$, a non-lethal but intolerable tumor size if $N_{tol} < K_r/\beta < N_{crit}$, and eventual death if $K_r/\beta > N_{crit}$.[13]

If $\beta$ is very high, that is, if sensitive cells strongly inhibit the growth of resistant cells, then it could be that $K_r/\beta < N_{tol}$ though $K_r > N_{crit}$. In that case, an aggressive treatment would lead to death, even though a containment strategy would have allowed stabilizing tumor size at a tolerable level forever.

Of course, any conclusion that under some circumstances containment could last forever is dubious, as our model then loses validity. If containment is expected to last for a very long time, a model that better accounts for the long-term evolution of the tumor should be developed, taking into account the appearance of new cell phenotypes.

5.2. **Approximate formula for the benefit of containment with resistance costs.** We derive here an approximate formula for the relative benefit of ideal containment versus ideal MTD in terms of time to treatment failure:

$$\frac{t_{fail}(idContN_{tol})}{t_{fail}(idMTD)},$$

for the above model (Model 3 in the main text). This formula is used to plot Fig. 4a in the main text. The approximation is valid if resistant cells are initially very rare and do not grow much more quickly than sensitive cells in the absence of treatment. Since ideal MTD instantly eliminates sensitive cells, resistant costs introduce no difficulty. The time to treatment failure is given by (see Section 3):

$$t_{fail}(idMTD) = \frac{1}{\rho_r} \ln\left(\frac{\ln(K_r/R_0)}{\ln(K_r/N_{tol})}\right)$$

(unless $K_r \leq N_{tol}$, that is, $\gamma \geq K_s/N_{tol} \simeq 28.6$, in which case a resistant tumor is benign, always tolerable, and time to treatment failure is infinite).

To compute the time to treatment failure of ideal containment, we distinguish two phases: the growth until $N_{tol}$, and the stabilization phase. Consider the first phase. Assuming that the initial frequency of resistant cells is very small, and that they do not expand much faster than sensitive cells, then the tumor remains almost fully sensitive during this phase. The dynamics of the total population $N$ may then be

[13] Favourable outcomes need not be attainable if the initial tumor is too resistant.

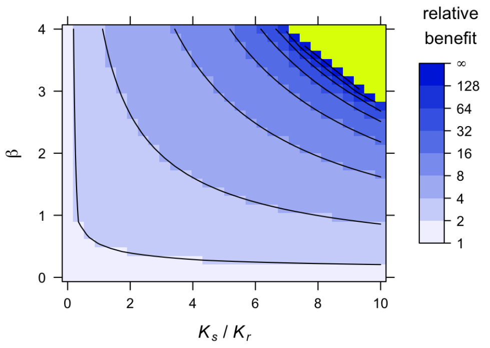


Supplementary figure 5. **Consequences of costs of resistance in a Gompertzian growth model (Model 3 in the main text).** Relative benefit, in terms of time to treatment failure, for ideal containment (at size $N_{tol}$) versus ideal MTD, for varied values of $K_r$ and $\beta$. The figure is obtained from simulations, while Fig. 4a in the main text is obtained from our approximate formula. Contour lines are at powers of 2. Fixed parameter values are as in Table 1 of the main text.

approximated by the dynamics of a fully sensitive tumor:

$$\dot{N} = \rho_s N \ln(K_s/N).$$

It follows that the time it takes for the tumor to grow from $N_0$ to $N_{tol}$ is approximately:

$$t_{N_0 \to N_{tol}} \simeq \frac{1}{\rho_s} \ln\left(\frac{\ln(K_s/N_0)}{\ln(K_s/N_{tol})}\right). \tag{13}$$

Moreover, during this phase (with the approximation $S = N$, $R = 0$, to compute the per-cell growth rates),

$$\frac{1}{\rho_r}\frac{\dot{R}}{R} - \frac{1}{\rho_s}\frac{\dot{S}}{S} \simeq -\ln(K_s/K_r).$$

A new approximation $S = N$ leads to:

$$R \simeq R_0 \left(\frac{N}{N_0}\right)^{\rho_r/\rho_s} \exp(-t\rho_r \ln(\beta K_s/K_r))$$

so that, letting $R_1$ denote the resistant population size at the beginning of the stabilization phase:

$$R_1 \simeq R_0 \left(\frac{N_{tol}}{N_0}\right)^{\rho_r/\rho_s} \exp(-t_{N_0 \to N_{tol}} \rho_r \ln(\beta K_s/K_r)).$$

That is,

$$R_1 \simeq R_0 \left(\frac{N_{tol}}{N_0}\right)^{\rho_r/\rho_s} \exp\left(-\frac{\rho_r}{\rho_s} \ln\left(\frac{\ln(K_s/N_0)}{\ln(K_s/N_{tol})}\right) \ln(\beta\gamma)\right) \tag{14}$$

with $\gamma = K_s/K_r$. Finally, the length $t_{stab}$ of the stabilization phase depends on whether the resistant population grows or not when the tumor reaches $N_{tol}$. If it decreases (which is the case if $(1-\beta)R_1 + \beta N_{tol} \geq K_r$, which essentially boils down to $\beta\gamma > K_s/N_{tol} \simeq 28,6$ for initially rare resistant cells), then the treatment never fails:

$$t_{stab} = +\infty.$$

Otherwise,

$$t_{stab} = \frac{1}{\rho_r} \int_{R_1}^{N_{tol}} \frac{dR}{Rg(R)} \text{ with } g(R) = \ln\left(\frac{K_r}{R + \beta(N_{tol} - R)}\right) \tag{15}$$

and $R_1$ is given approximately by (14). Summing the terms in (13) and (15) leads to an approximation of time to treatment failure under ideal containment at $N_{tol}$ since:

$$t_{fail}(idContNtol) = t_{N_0 \to N_{tol}} + t_{stab}.$$

The formula is not precise for very low values of $\beta$, because resistant cells then grow much more quickly than sensitive cells as long as they are rare, so that the approximation of an almost fully sensitive tumor during the growth phase from $N_0$ to $N_{tol}$ need not be appropriate. The comparison of Fig. 4a and Supplementary Fig. 5 shows that the figure obtained from this approximate formula in a Gompertzian growth model is similar to figures obtained by simulations, which require heavy computation.

*Remark:* Integrating from $R_1$ to $R_2$ in (15), where $R_2$ is the resistant population size when containment fails (see Section 3.1.4) leads to a formula for time to treatment failure under containment (as opposed to ideal containment). Moreover, the same method (approximating tumor growth by the growth of a fully sensitive tumor in the initial phase) allows one to find approximate formulas for time to treatment failure for more general models.

**The case $\beta = 1$.** For $\beta \neq 1$, we were unable to solve explicitly the integral in (15) and had to evaluate it numerically. But if $\beta = 1$, computing this integral is easy, since $g(R)$ is then constant. This leads to:

$$(\text{Case } \beta = 1) \quad t_{stab} = \frac{1}{\rho_r} \frac{\ln(N_{tol}/R_1)}{\ln(K_r/N_{tol})}.$$

We then obtain an explicit approximate formula for the time to treatment failure under ideal containment. This formula is simpler when $\rho_r = \rho_s = 1$. Letting

$$A = \ln\left(\frac{\ln(K_s/N_0)}{\ln(K_s/N_{tol})}\right),$$

we then find:

$$(\text{Case } \beta = 1,\ \rho_r = \rho_s = 1) \quad t_{fail}(idCont) \simeq A + \frac{\ln(N_0/R_0) + A\ln(\gamma)}{\ln(K_r/N_{tol})}.$$

The ratio is then approximately:

$$(\text{Case } \beta = 1, \rho_r = \rho_s = 1) \quad \frac{t_{fail}(idCont)}{t_{fail}(idMTD)} \simeq \frac{A + \frac{\ln(N_0/R_0) + A\ln(\gamma)}{\ln(K_r/N_{tol})}}{\ln\left(\frac{\ln(K_r/R_0)}{\ln(K_r/N_{tol})}\right)}.$$

#### 6. Containment at the initial size in practice

One of our findings is that, at least for our simple models, the precise way to implement containment treatments is not essential. Nevertheless, we discuss here how containment at the initial tumor size $N_0$ could be attempted in practice: how should the initial dose be chosen? how to adapt the dose during the course of treatment ? how many measurements are likely to be needed before finding a dose that approximately stabilizes tumor size? We assume that some biomarker allows us to estimate tumor size (or its variation). This biomarker level is measured at times $t_0$, $t_1$,...,$t_k$,..., leading to an estimated tumor size of $N_k$ at time $t_k$ (or an estimated ratio $N_k/N_0$). A constant dose $C_k$ is given between times $t_k$ and $t_{k+1}$. Though the timing of the measurements is important, we focus on the choice of the doses $C_k$.[14]

##### 6.1. Some simple protocols.

We first discuss some simple protocols, before proposing a slightly more elaborated one.

*Downward titration*: start with the maximal tolerated dose, and then decrease the dose by a certain fraction of the maximal tolerated dose until tumor size (or rather, the biomarker level) starts increasing. E.g. use initially the dose $C = C_{max}$, then $C = 0.9\,C_{max}$, then $C = 0.8\,C_{max}$ and so on, until the dose is low enough for tumor size to increase again. At that point, the dose would remain constant until tumor size becomes higher than at the beginning of treatment (or could be decreased again if for some reason tumor size starts decreasing while it is still below its initial level $N_0$). Once tumor size becomes higher than $N_0$, the dose would be incremented upwards until tumor size starts decreasing, and so on. A variant is to use increments that are not a certain fraction of $C_{max}$ but of the current dose. For instance, if the dose $C_k$ at step $k$ is too weak, the next dose could be $C_{k+1} = 1.1C_k$, instead of $C_{k+1} = C_k + 0.1C_{max}$.

*Upward titration*: similar, but starting with a low dose instead of a high dose, and gradually increasing it until a dose is found that allows to decrease tumor size.

*The Gallaher et al. (2018) protocol*: Jill Gallaher and collaborators consider a protocol that is essentially a downward titration method but with some twists. First, a treatment vacation occurs if tumor size becomes smaller than half the initial size. Second, the dose does not change if tumor size varied little since

[14] It makes sense to have shorter time intervals between the first measurements, in order to assess patient's specific reaction to treatment, and also when recent measurements suggest a quick evolution of tumor size, or measurements errors.

the last measurement. With our notation, the protocol may be described as follows: initially, $C_0 = C_{max}$. Later, if $N_{k+1} < N_0/2$, a treatment vacation occurs: $C_{k+1} = 0$. Otherwise,

$$C_{k+1} = \begin{cases} \min(C_{max}, (1+\alpha)C_k^*) & \text{if } N_{k+1}/N_k > 1+\beta \\ C_k^* & \text{if } 1-\beta < N_{k+1}/N_k \leq 1+\beta \\ (1-\alpha)C_k & \text{if } N_{k+1}/N_k \leq (1-\beta) \end{cases}$$

where $\alpha$ and $\beta$ are parameters (e.g., $\alpha = 0.25$, $\beta = 0.05$), and $C_k^* = C_k$ unless $C_k = 0$, in which case $C_k^*$ denotes the last positive dose given. In other words, if a treatment vacation occurs, the reference dose to determine further modulations does not become zero but remains equal to the last positive dose.[15]

Titration methods have the advantage of being conceptually simple. However, they might be slow in determining an approximately stabilizing dose. For downward titration, this could result in too strong an initial treatment, and competitive release. For upward titration, this could result in the tumor growing very large before treatment is increased sufficiently to stabilize it.

Moreover, the methods we described do not fully take into account how far the current tumor size is from its target, and how much tumor size decreased or increased since the last measurement. For instance, assume that in the Gallaher et al. protocol, $\alpha = 0.25$ and $\beta = 0.05$, so that the dose is increased by 25% if tumor size increased by more than 5%, but does not change if tumor size changed by less than 5% since the last measurement. Except when a treatment vacation occurs, this rule does not depend on whether tumor size if close or far from some target[16]. More importantly, the modulation rule does not differentiate between a tumor that increased by 4% and a tumor that decreased by the same amount, nor does it differentiate between a tumor that increased by 6% and a tumor that increased by 60%. Finally, Gallaher et al.'s protocol is sensitive to the time interval between two measurements and the speed of evolution of the tumor, which may be patient specific: if measurements are frequent, tumor evolution is slow, or $\beta$ is large, then it may be that tumor size never evolves sufficiently between two successive measurements to trigger dose modulation, though in the long run tumor size may evolve substantially.

We now propose some ideas to improve these methods. First, starting from an intermediate dose rather than from a very high or very low dose should somewhat speed up the process of determining a stabilizing dose. How large the initial dose should be depends on how large the tumor currently is, that is, whether the main issue is to quickly avoid tumor size growing larger, or to avoid triggering competitive release.

Second, if, e.g., the first dose is too high, then instead of slightly decreasing it again and again until the tumor starts to rebounce, we suggest to decrease it relatively sharply. It is then more likely that the second dose will be lower than the stabilizing dose, helping tumor size not to drift too far away from its target. Large initial adjustments also help to adapt the treatment sufficiently quickly when patient's reaction is untypical (e.g., the patient is particularly responsive or unresponsive). How sharply the dose should be changed between the first and the second dose would depend on whether tumor size evolved a lot or just a little between the first two measurements.

Third, once the effect of the first two doses is observed, that is, after three measurements, an educated guess for a stabilizing dose could be achieved by building on simple models of tumor growth. This is explained below in Section 6.3. Based on the above considerations, we now propose a protocol taking into account how far tumor size is from target and how much it recently increased or decreased.

6.2. **A new protocol.** A basic family of protocols is as follows (see remarks afterwards for details and refinements). The first two doses are somewhat arbitrary.

1. At time $t_0$: choose an initial dose expected to at least stabilize tumor size in most similar patients (e.g. in 75% of similar patients). [17]

2. At time $t_1$: given patient's response to the first dose, choose a dose expected to bring tumor size back towards $N_0$.

[15] The description in the original article is somewhat different, but Jill Gallaher kindly confirmed that what was meant and implemented is the above described protocol.

[16] This need not be a defect, it depends whether the emphasis is on stabilizing the tumor at any level, or on stabilizing it close to its initial size.

[17] The choice of 75% of patients is arbitrary, but at least in initial clinical trials, patients and physicians are likely to feel more comfortable with not too low an initial dose. In theory, whether choosing a first dose that would work for an average patient, or for a large majority of patients, depends on whether current tumor size is considered worrying large or not. If the initial tumor size seems way below our hypothetical maximal tolerable size, then starting with a relatively low dose is reasonable. If the initial tumor size is quite large, then starting with a relatively high dose is bound to be preferable.

3. From time $t_2$ on: compute an estimation $C_{guess}$ of the dose that would currently stabilize the tumor, based on recent measurements and a standard tumor growth model, e.g., using formulas of Section 6.3 below. Deliver this dose if $N = N_0$, a higher dose if $N > N_0$, and a lower one if $N < N_0$.

As mentioned in the main text, a concrete example is to fix a low and a high threshold, $N_l < N_0$ and $N_h > N_0$, positive parameters $\gamma_1$ and $\gamma_2$, and to deliver the following dose between time $t_k$ and $t_{k+1}$:

$$C_k = \begin{cases} 0 & \text{if } N_k \leq N_l; \\ C_{guess}\left[1 - \left(\frac{N_0 - N}{N_0 - N_l}\right)^{\gamma_1}\right] & \text{if } N_l \leq N_k \leq N_0; \\ C_{guess} + (C_{max} - C_{guess})\left(\frac{N - N_0}{N_h - N_0}\right)^{\gamma_2} & \text{if } N_0 \leq N_k \leq N_h; \\ C_{max} & \text{if } N_k \geq N_h \end{cases}$$

Thus, the dose given depends both on the estimated stabilizing dose and on how far tumor size is to its target $N_0$. A treatment vacation occurs if tumor size falls below the low threshold, and the tumor is treated at the maximal tolerated dose if tumor size increases beyond the high threshold. In between, the dose is a continuously increasing function of tumor size, equal to the estimated stabilizing dose if tumor size is equal to its target $N_0$. The parameters $\gamma_1$ and $\gamma_2$ tune whether the emphasis is on stabilizing tumor size ($\gamma_i > 1$) or bringing it back to $N_0$ ($\gamma_i < 1$). For instance, if $\gamma_2$ is significantly larger than 1, then it is only when tumor size approaches the higher threshold $N_h$ that the dose given becomes substantially different from the estimated stabilizing dose.

Some remarks are in order:

a) The above protocol is for containment at the initial size, but may easily be adapted for containment at another target size.

b) A target size, whether absolute or relative to initial tumor size, may prove too large for some patients, in the sense of leading to a low quality of life or other adverse effects. To deal with this issue, the target size could be defined as the minimum of an a priori target size (e.g., the initial size) and the largest tumor size compatible with a satisfying quality of life.

c) Protocols should be adapted to each tumor type and biomarker. If the link between tumor size and biomarker level is known to evolve with time, the protocol should be modified accordingly. [18]

d) If containment treatments become common, data on previous patients should allow to determine the percentage of tumors stabilized by a given dose. This would help choosing the initial dose. For initial clinical trials, an educated guess could be made using the ideas of Section 6.3, provided tumor growth data is available in the absence of treatment and under standard of care.

e) If tumor growth data in the absence of treatment is available for the current patient, then an educated guess of the stabilizing dose for this specific patient could be computed already for the second dose (at time $t_1$).

f) The second measurement should be made shortly after the first one, since the first dose might be well off the mark due to patients' heterogeneity. The third measurement should also be made quickly if the second one revealed quick evolution of tumor size. Once an approximately stabilizing dose has been found, tumor should evolve more slowly, and monitoring could become rarer. However, tumor's response to treatment is likely to evolve with time. Thus, regular monitoring remains needed.

g) Tumor's true response, measurements of biomarkers, and the relation between tumor size and biomarkers are likely to be noisy. In order not to rely excessively on a very small number of possibly wrong data points, and to smooth out the noise, methods could be developed to base the estimate of the current stabilizing dose not only on the last three data points but on all available data points. These methods should nevertheless give a greater weight to recent data, to take into account tumor evolution. Here is a simple possibility. Let $C_{guess}(t_k)$ denote the stabilizing dose between $t_k$ and $t_{k+1}$ suggested by the last three data points (from times $t_{k-2}$, $t_{k-1}$ and $t_k$). Define the smoothed stabilizing dose by $C_{smooth}(t_2) = C_{guess}(t_2)$, and for $k \geq 3$,

$$C_{smooth}(t_k) = (1 - \delta)C_{guess}(t_k) + \delta C_{smooth}(t_{k-1}),$$

where $\delta \in (0, 1)$ is a parameter tuning the importance given to recent data compared to older data. More involved methods with an explicit modeling of the noise could also be considered.

[18] In Zhang et al.'s (2017) clinical trial of adaptive therapy on metastatic castrate resistance prostate cancer, the biomarker used is prostate specific antigen (PSA). An issue is that cells resistant to the drug used, Abiraterone, may contribute much more to PSA production than sensitive cells, thus an increase in PSA might signal an increase in tumor size or an increase in the fraction of resistant cells.

h) Different models might give different estimates of the current stabilizing dose. To eventually rely on the best models, a possibility is to use as our estimates for the stabilizing dose a weighted average of estimates produced by a number of different models, with larger weights put on models that fit well previous data points. There is a large literature in statistics and game theory on how to choose the best "expert", in our case, the best model [23].

Complimentary considerations on practical implementation of containment treatments may be found in the forthcoming Ph.D. dissertation of Jessica Cunnhingham.

6.3. **How to make an educated guess for the current stabilizing dose?** We conclude by explaining how to make an educated guess for the stabilizing dose based on the last three measurements. Assume to begin with that a tumor growth-model of the form

$$\dot{N} = Ng(N)(1 - \lambda C) \tag{16}$$

is deemed reasonable. Assume also that between $t_{k-2}$ and $t_k$, tumor size does not evolve too much so that the natural net growth-rate $g(N)$ can be considered approximately equal to some constant $g$. Denote by $\rho_{k-2} = \frac{1}{t_{k-1}-t_{k-2}} \ln(N_{k-1}/N_{k-2})$ and $\rho_{k-1} = \frac{1}{t_k - t_{k-1}} \ln(N_k/N_{k-1})$ the average per-cell growth-rate on the time intervals $[t_0, t_1]$ and $[t_1, t_2]$, respectively. This leads to:

$$\begin{cases} \rho_{k-2} = g(1 - \lambda C_{k-2}) \\ \rho_{k-1} = g(1 - \lambda C_{k-1}) \end{cases}$$

Solving this system and noting that for Model (16) the stabilizing dose is $C = 1/\lambda$ leads to:[19]

$$C_{guess} = \frac{\rho_{k-1}C_{k-2} - \rho_{k-2}C_{k-1}}{\rho_{k-1} - \rho_{k-2}} \tag{17}$$

If the kill-rate is not assumed proportional to the dose but to some function of the dose:

$$\dot{N} = Ng(N)(1 - \lambda f(C))$$

for instance due to some saturation effect, then the estimated stabilizing dose is such that:

$$f(C_{guess}) = \frac{\rho_{k-1}f(C_{k-2}) - \rho_{k-2}f(C_{k-1})}{\rho_{k-1} - \rho_{k-2}} \tag{18}$$

and can be found by inverting the function $f$. This formula may also be expressed in terms of doubling times: letting $DT_{k-2} = (\ln 2)/\rho_{k-2}$ and $DT_{k-1} = (\ln 2)/\rho_{k-1}$ denote the tumor's doubling time on the time-intervals $[t_{k-2}, t_{k-1}]$ and $[t_{k-1}, t_k]$ leads to:

$$f(C_{guess}) = \frac{DT_{k-1}f(C_{k-1}) - DT_{k-2}f(C_{k-2})}{DT_{k-1} - DT_{k-2}}$$

The important point is that the exact same formulas are obtained for a variety of other models. For instance, if we use a log-kill rate instead of a Norton-Simon kill rate:

$$\dot{N} = N[g(N) - \tilde{\lambda} f(C)]$$

then we still obtain (18). This remains true with birth-death models, with a Norton-Simon kill rate

$$\dot{N} = N[b(N)(1 - \lambda f(C) - d(N)]$$

or a log-kill rate

$$\dot{N} = N[b(N) - d(N) - \tilde{\lambda} f(C)]$$

under the assumption that the birth and death rates $b(N)$, $d(N)$ are approximately constant.This is because, under these simplifying assumptions, the above four class of models are equivalent. Variants of these models with sensitive and resistant tumor cells, e.g., Model 2, also lead to the same formulas if the frequencies of each cell type may be considered constant between $t_{k-2}$ and $t_k$. Though we do not expect them to work perfectly, we conclude that the above formulas could provide reasonable initial guesses for stabilizing doses.[20]

---

[19] Note that only relative variations matter, that is, quotients $N_{k+1}/N_k$. This is handy when the evolution of biomarker level is correlated to the evolution of tumor size but the initial biomarker level is not much informative in itself, as is the case for prostate specific antigen level in prostate cancer.

[20] Due to measurement errors or some unexpected phenomenon, it might be that though the dose was recently increased: $C_{k-1} > C_{k-2}$, the estimated tumor growth rate also increased: $\rho_{k-1} > \rho_{k-2}$. This is incompatible with our deterministic models and the assumption that the frequency of resistant cells may be considered constant on $[t_{k-2}, t_k]$. The above formulas

#### 7. Taking mutations into account

In this section, we justify our choice of neglecting mutations from sensitive to resistant cells. To do so, consider the following generalization of Monro and Gaffney's (2009) [11] model:

$$\begin{aligned} \dot{S} &= g(N)[(1-\tau_1)S + \tau_2 R] - d(C,N)S \\ \dot{R} &= g(N)[(1-\tau_2)R + \tau_1 S] \end{aligned} \tag{19}$$

The constants $\tau_1$ and $\tau_2$ are mutations and back-mutations rates, and $d(C,N)$ is a treatment induced death rate that vanishes if $C = 0$. Note that the number of mutations is assumed proportional to the net growth rate, so if the death to birth ratio is high, the above "mutation rates" could be much higher than true mutations rates (proportional to the birth rates). For these reasons, high values of $\tau_1$ and $\tau_2$ will be considered. A variant where mutations are explicitly proportional to birth rates is studied in Variant 2 below.

Let $t_{mut}$ denote time to progression under containment at the initial size $N_0$. Let $R_{prog}$ be the size of the resistant population at progression. During the stabilization phase, $N = N_0$, hence

$$\dot{R} = g(N_0)[(1-\tau_2)R + \tau_1(N_0 - R)] = g(N_0)[aR + b]$$

with $a = 1 - \tau_1 - \tau_2$, and $b = \tau_1 N_0$. Solving this linear differential equation leads to:

$$t_{mut} = \frac{1}{ag(N_0)} \ln\left(\frac{R_{prog} + b/a}{R_0 + b/a}\right) = \frac{1}{(1-\tau_1-\tau_2)g(N_0)} \ln\left(\frac{f_{prog} + \tau_1/(1-\tau_1-\tau_2)}{f_0 + \tau_1/(1-\tau_1-\tau_2)}\right) \tag{20}$$

where $f_{prog} = R_{prog}/N_0$ and $f_0 = R_0/N_0$ are the frequencies of resistance at progression and at treatment initiation, respectively. The time to progression $t_{nomut}$ for the model neglecting mutations is recovered by letting $\tau_1 = \tau_2 = 0$.

Assuming that the tumor is initiated with $S = 1$ and $R = 0$, the number of resistant cells for a tumor of size $N$ is (Goldie and Coldman, 1979 [20], Luria and Delbrück, 1943 [24]),

$$R(N) = \frac{\tau_1}{\tau_1 + \tau_2} N(1 - N^{-\tau_1 - \tau_2}) \simeq \tau_1 N \ln N \tag{21}$$

for $\tau_1 + \tau_2$ small. This leads to:

$$\frac{t_{nomut} - t_{mut}}{t_{nomut}} \simeq \frac{\ln\left(1 + \frac{1}{\ln N_0}\right)}{\ln(f_{prog}/\tau_1 \ln N_0)}$$

Assuming $N_0 = 10^{10}$, $R_0 = R(N_0)$ (as defined in (21)) and either $\tau_1 = \tau_2$ as in [11], or $\tau_2 = 0$ (no backmutations), Table 10 gives the relative difference $(t_{nomut} - t_{mut})/t_{nomut}$ for various values of $\tau_1$ and $f_{prog}$ ($f_{prog} = 1$ corresponds to ideal containment). This is based on exact formulas, not the above approximation.

We find that the impact of ongoing mutations on time to progression is typically less than 1%. This is much less than the gain of containment strategies over aggressive strategies that we found in our simulations. This would also be true of time to treatment failure or survival time. This suggests that, for a Gompertz model, taking into account mutations after treatment initiation has little impact on the comparison between containment and aggressive strategies. To test the robustness of this finding, we examine three variants. These three variants all lead to a smaller initial resistant population than in (21), due to late appearance of the first resistant cell (Variant 1), an increasing death to birth ratio (Variant 2), or slow replication of resistant cells (Variant 3).

then should not be used directly, though a smoothened version of the estimated stabilizing dose, as discussed in Section 6.2, could still be a useful indicator.

Supplementary table 10. **Impact of ongoing mutations on time to progression of ideal containment in a general density-dependent model.** Ratio $(t_{nomut} - t_{mut})/t_{nomut}$ in Model (19), for various mutations rates and frequency of resistant cells at progression, with $N_0 = 10^{10}$, and either $\tau_2 = \tau_1$ or, in parenthesis, $\tau_2 = 0$.

| $\tau_1$ | $10^{-6}$ | $10^{-5}$ | $10^{-4}$ | $10^{-3}$ |
|---|---|---|---|---|
| $f_{prog} = 1$ | 0.40% (0.40%) | 0.51% (0.51%) | 0.68% (0.69%) | 0.92% (1.01%) |
| $f_{prog} = 1/2$ | 0.43% (0.43%) | 0.55% (0.55%) | 0.77% (0.77%) | 1.14% (1.22%) |

Supplementary table 11. **Impact of ongoing mutations on time to progression of ideal containment if resistant cells appear late.** The table gives the ratio $(t_{nomut} - t_{mut})/t_{nomut}$ in Model (19), with late initial mutation (see text), treatment initiated at $N_0 = 10^{10}$, and $\tau_2 = \tau_1$.

| $\tau_1$ | $10^{-6}$ | $10^{-5}$ | $10^{-4}$ | $10^{-3}$ |
|---|---|---|---|---|
| $f_{prog} = 1$ | 1.00% | 1.02% | 1.13% | 1.34% |
| $f_{prog} = 1/2$ | 1.08% | 1.10% | 1.26% | 1.62% |

7.1. **Variant 1: Late first mutation.** Mutations are a stochastic phenomenon, so the resistant population could be – and is actually likely to be – smaller than the above estimate (Goldie and Coldman, 1979 [20]). To get an idea of what would happen then, let $R_0^*$ denote the resistant population when $N = 10^{10}$ found by solving (19) with initial condition $R = 1$ and $S = S^*$, where $S^* = -2(\ln 10)/\ln(1-\tau_1) \simeq 4,6/\tau_1$. This is the value such that there is a 99% probability that the first resistant mutant appears when $S \leq S^*$. Table 11 gives the relative difference $(t_{nomut} - t_{mut})/t_{nomut}$ for $N_0 = 10^{10}$, and $R_0 = R_0^*$.

The conclusion is that even if the first resistant cell appears late, taking into account mutations after treatment initiation would not diminish time to progression under containment or ideal containment by more than 1 or 2%. This is still much less than the gain of containment strategies over aggressive strategies that we found in our simulations.

7.2. **Variant 2: Birth-death model.** As mentioned before, in Model (19), the number of mutations is proportional to the net growth rate rather than the birth rate. To deal with this issue, consider the following birth-death model:

$$\begin{aligned} \dot{S} &= b(N)[(1-\tau_1)S + \tau_2 R] - d(N)S - d_{tr}(C,N)S \\ \dot{R} &= b(N)[(1-\tau_2)R + \tau_1 S] - d(N)R \end{aligned} \tag{22}$$

where $b(N)$ is the birth rate, $d(N)$ a death rate common to both types, and $d_{tr}(N,C)$ a treatment-induced additional death rate that vanishes when $C = 0$. Let $g(N) = b(N) - d(N)$ denote the natural net growth rate and let $q(N) = b(N)/g(N)$. The above equations may be rewritten as:

$$\begin{aligned} \dot{S} &= g(N)[(1-q(N)\tau_1)S + q(N)\tau_2 R] - d_{tr}(C,N)S \\ \dot{R} &= g(N)[(1-q(N)\tau_2)R + q(N)\tau_1 S] \end{aligned} \tag{23}$$

If the death to birth ratio is constant, then so is $q(N)$: $q(N) = q$. We then obtain the same model as in (19), but with mutation rates increased by a factor $q$. Taking mutations into account would then have a very small impact.

If the death to birth ratio is increasing, as one would expect (Monro and Gaffney, 2009, Appendix A.4 [11]), then before treatment initiation $q(1) \leq q(N) \leq q(N_0)$. It follows from (23) that:

$$\frac{dR}{dN} = \frac{R}{N} + q(N)\left[\tau_1 - \frac{(\tau_1+\tau_2)R}{N}\right]$$

Unless $\tau_2 \gg \tau_1$, the bracket should be positive before treatment initiation, so that $dR/dN$ is increasing in $q(N)$. Comparison principles (variants of Gronwall's lemma, see Section 2, Property 10) then lead to:

$$R(N_0) = \frac{\tau_1}{\tau_1+\tau_2} N_0 (1 - N_0^{-\bar{q}(\tau_1+\tau_2)}) \simeq \bar{q}\tau_1 N_0 \ln N_0 \tag{24}$$

for some $\bar{q}$ in $(q(1), q(N_0))$. The time to progression $t_{mut}$ is obtained from (20) by replacing $\tau_1$ and $\tau_2$ by $q(N_0)\tau_1$ and $q(N_0)\tau_2$, and letting $R_0 = R(N_0)$ as defined in (24).

Assuming $N_0 = 10^{10}$, $\tau_1 = \tau_2$, and $q(N_0) = 10$ (which corresponds to a death to birth ratio $d(N_0)/b(N_0) = 9/10$), Table 12 gives the relative difference of times to progression: $(t_{nomut} - t_{mut})/t_{nomut}$ for various values of $\tau_1$ and of the ratio $\bar{q}/q(N_0)$.

The impact of taking into account mutations after treatment initiation is stronger when $q(N)$ (or equivalently the death to birth ratio) is much higher at treatment initiation than during the initial development of the tumor. However, the effect remains limited, especially if we take into account that high mutations rates are included above only for consistency between tables: they were considered previously as a shortcut for such an explicit birth-death model with a high death to birth ratio, and need not be meaningful for the current model.

Supplementary table 12. **Impact of ongoing mutations on time to progression of ideal containment in the birth-death model** (22)**.** The table gives the ratio $(t_{nomut} - t_{mut})/t_{nomut}$ with $N_0 = 10^{10}$, $\tau_2 = \tau_1$, and $\bar{q}$ defined by (24).

| $\tau_1$ | $10^{-6}$ | $10^{-5}$ | $10^{-4}$ | $10^{-3}$ |
|---|---|---|---|---|
| $\bar{q} = q(N_0)$ | 0.51% | 0.68% | 0.92% | 0.60% |
| $\bar{q} = q(N_0)/2$ | 0.91% | 1.21% | 1.67% | 1.75% |
| $\bar{q} = q(N_0)/5$ | 1.97% | 2.54% | 3.46% | 4.44% |

7.3. **Variant 3: Slowly growing resistant cells.** The last variant we consider is to assume that the resistant cells expand much slower than sensitive cells. In this case, a large part of the resistant population growth at the time of treatment initiation should come from mutations from sensitive to resistant cells, and taking into account mutations should have a larger impact. To examine this issue formally, consider the following variant of Eq. (19), with type-specific growth rates.

$$\begin{aligned} \dot{S} &= g(N)[(1-\tau_1)\rho_s S + \tau_2 \rho_r R] - d(C,N)S \\ \dot{R} &= g(N)[(1-\tau_2)\rho_r R + \tau_1 \rho_s S] \end{aligned} \tag{25}$$

Eq. (20) is then replaced by:

$$t_{mut} = \frac{1}{[(1-\tau_2)\rho_r - \tau_1\rho_s]g(N_0)} \ln\left(\frac{f_{prog} + \tau_1\rho_s/[(1-\tau_2)\rho_r - \tau_1\rho_s]}{f_0 + \tau_1\rho_s/[(1-\tau_2)\rho_r - \tau_1\rho_s]}\right) \tag{26}$$

To estimate the initial frequency of resistant cells, note that if we neglect back mutations (a reasonable approximation for $\tau_1 \leq 10^{-3}$) then we obtain the linear differential equation:

$$\frac{dR}{dS} = \frac{\tilde{a}}{S}R + \tilde{b}, \text{ with } \tilde{a} = \frac{\rho_r}{\rho_s(1-\tau_1)}, \; \tilde{b} = \frac{\tau_1}{1-\tau_1}.$$

Assuming that $R = 0$ when $S = 1$ leads to

$$R(S) = \frac{\tilde{b}}{1-\tilde{a}} S\left(1 - S^{-(1-\tilde{a})}\right) = \frac{\tau_1}{1-\tau_1-\rho_r/\rho_s}\left[S - S^{\frac{\rho_r}{\rho_s(1-\tau_1)}}\right] \tag{27}$$

If $\rho_r$ is substantially smaller than $\rho_s$, then for $S$ large, $S^{\rho_r/\rho_s(1-\tau_1)} \ll S$, and $1 - \tau_1 - \rho_r/\rho_s \simeq 1 - \rho_r/\rho_s$. This leads to:

$$R(s) \simeq \frac{\tau_1 S}{1-\rho_r/\rho_s} \simeq \frac{\tau_1 N}{1-\rho_r/\rho_s}$$

and a relative difference

$$\frac{t_{nomut} - t_{mut}}{t_{nomut}} \simeq \frac{\ln(\rho_s/\rho_r)}{\ln(1-\rho_r/\rho_s) - \ln(\tau_1)}$$

Table 13 gives the estimate of the relative difference between times to progression of ideal containment with and without mutations: $(t_{nomut} - t_{mut})/t_{nomut}$, found by using Equations (26) and (27) for a treatment beginning when $S = 10^{10}$.[21]

This shows that if $\rho_r$ is much smaller than $\rho_s$, then taking into account mutations could decrease substantially time to progression under containment strategies. Note however that a ratio $\rho_r/\rho_s = 0.5$

[21]Initiating treatment when $N = 10^{10}$ or neglecting back mutations in (26) would change these values very little.

Supplementary table 13. **Impact of ongoing mutations on time to progression of ideal containment with slowly growing resistant cells.** The table gives the ratio $(t_{nomut} - t_{mut})/t_{nomut}$ in Model (25) for various mutations rates and baseline growth rates of resistant cells, with $N_0 = 10^{10}$, $\tau_2 = \tau_1$.

| $\tau_1$ | $10^{-6}$ | $10^{-5}$ | $10^{-4}$ | $10^{-3}$ |
|---|---|---|---|---|
| $\rho_r/\rho_s = 1$ | 0.40% | 0.51% | 0.68% | 0.91% |
| $\rho_r/\rho_s = 0.8$ | 1.84% | 2.27% | 2.94% | 4.01% |
| $\rho_r/\rho_s = 0.5$ | 5.28% | 6.40% | 8.11% | 10.88% |
| $\rho_r/\rho_s = 0.2$ | 11.84% | 14.26% | 17.86% | 23.62% |

would already be a huge resistance cost. Moreover, this does not imply that, in a model with mutations, reducing $\rho_r$ would decrease the relative gain of ideal containment over ideal MTD.

To explore this issue, let $t_{idMTD}$ and $t_{mut}$ denote times to progression of ideal MTD and of ideal containment in a model with mutation terms. As the two other variants we considered, reducing $\rho_r$ leads to a smaller initial resistant population than in (21). On the one hand, this tends to increase the proportion of the growth of the resistant population that comes from mutations from sensitive to resistant cells: this decreases $t_{mut}/t_{nomut}$. On the other hand, this increases the per-cell growth rate of the tumor if sensitive cells are eliminated: this decreases $t_{idMTD}/t_{nomut}$. The net effect on the ratio $t_{idMTD}/t_{mut}$ is unclear.

Consider for instance the Gompertz model of Monro and Gaffney that we used in simulations, that is, with $g(N) = \ln(K/N)$, $K = 2 \times 10^{12}$ and $N_0 = 10^{10}$. Table 14 gives the relative gain of ideal containment over ideal MTD: $(t_{mut} - t_{idMTD})/t_{mut}$, for various values of $\tau_1$ and $\rho_r/\rho_s$. It shows that, for this model, the relative gain of ideal containment over ideal MTD initially increases when $\rho_r$ decreases, but then decreases when $\rho_r$ approaches 0. Though it increases the impact of mutations, a small resistance cost in the form of a reduction of $\rho_r$ thus increases the relative gain of ideal containment over ideal MTD.

7.4. **Discussion: links with Hansen et al. (2017).** Though seemingly contradictory, our analysis is consistent with the results of Hansen et al (2017) [8]. Extending the analysis of Martin et al. (1992) [6], they argue that if the initial resistant population is small enough, then eliminating sensitive cells is preferable to ideal containment. For a given mutation rate $\tau_1$, this is correct and consistent with Eq. (20).[22] However, our analysis suggests that this typically requires one of the two following conditions: either a tumor growth model leading to a very small advantage of ideal containment over ideal MTD when mutations are neglected: there is then no strong theoretical case for containment strategies anyway; or an abnormally small resistant population given the mutation rate $\tau_1$. With a Gompertz growth model and a resistant population size coherent with mutation rates, taking into account mutations from sensitive cells to resistant cells after treatment initiation does not seem to make containment strategies inferior to more aggressive ones. Note however that we only considered the case of random genetic mutations, independent of treatment. It would be important to examine the case of treatment-induced mutations or phenotypic changes.

[22] This must be, since if there were no resistant cells, eliminating sensitive cells would cure the tumor, and under ideal MTD, time to progression depends continuously on the initial proportion of resistant cells.

Supplementary table 14. **Relative gains of ideal containment over ideal MTD for Gompertzian growth with slowly growing resistant cells.** The table gives the ratio $(t_{mut} - t_{idMTD})/t_{mut}$ in Model (19) with $g(N) = \ln(K/N)$, $K = 2 \times 10^{12}$, $N_0 = 10^{10}$, $\tau_2 = \tau_1$, for various mutation rates and resistant baseline growth rates.

| $\tau_1$ | $10^{-6}$ | $10^{-5}$ | $10^{-4}$ | $10^{-3}$ |
|---|---|---|---|---|
| $\rho_r/\rho_s = 1$ | 45.02% | 39.72% | 32.93% | 23.90% |
| $\rho_r/\rho_s = 0.8$ | 47.17% | 42.32% | 36.13% | 27.85% |
| $\rho_r/\rho_s = 0.5$ | 46.88% | 41.79% | 35.12% | 25.77% |
| $\rho_r/\rho_s = 0.2$ | 43.79% | 37.54% | 28.81% | 15.32% |
